# Serial passage of the human probiotic *E. coli* Nissle 1917 in an insect host leads to changed bacterial phenotypes

**DOI:** 10.1101/2021.01.04.425298

**Authors:** Nicolas C. H. Schröder, Ana Korša, Haleluya Wami, Ulrich Dobrindt, Joachim Kurtz

**Affiliations:** Institute for Evolution and Biodiversity, WWU Münster, Germany; Institute for Hygiene, UKM Münster, Germany

## Abstract

Probiotics are living microorganisms that are increasingly and successfully used for the therapy of various diseases. The most common use of probiotics is the therapeutic and preventive application for gastrointestinal disorders. The probiotic *Escherichia coli* strain Nissle 1917 (*Ec*N) has been proven to effectively prevent and alleviate intestinal diseases, including various types of inflammatory bowel disease. Despite the widespread medical application of *Ec*N, the underlying mechanisms of its protective effect remain elusive. The present work aimed to establish an insect model system to enable further research on the modes of action of *Ec*N and the dynamics of adaptation to a novel host organism. Using a long-term serial passage approach, we orally introduced *Ec*N to the host, the red flour beetle *Tribolium castaneum*. After multiple cycles of intestinal colonization in beetle larvae, several attributes of the passaged replicate lines were assessed. We observed phenotypic changes in growth and motility but no genetic changes in the lines after passaging through the host and its flour environment. One of the *Ec*N lines exposed to the host displayed peculiar morphological and physiological characteristics showing that serial passage of *Ec*N can generate differential phenotypes.

## Introduction

Probiotics are the most prominent defensive microbes in humans. They are in use for thousands of years and studied for over a hundred years as therapeutics for dermal, vaginal and intestinal infections (Borges et al., 2014; Roudsari et al., 2015; Santacroce et al., 2019). Oral intake as a treatment against various gastrointestinal disorders, including severe bacterial infections, is the most widespread application of probiotics (Islam, 2016). As antibiotic resistance of pathogens might grow to a global public health problem with unforeseeable severity, the need for safe and well-characterized biotherapeutics asks for further research on the protective mechanisms of established probiotics (Imperial and Ibana, 2016; O’Toole et al., 2017).

A commonly used and medically important probiotic is the gram-negative bacterium *Escherichia coli* Nissle 1917 (*Ec*N). It belongs to the large family of *Enterobacteriaceae* and the *E. coli* serotype group O6:K5:H1 (Faubion and Sandborn, 2000). *Ec*N was introduced as the pharmaceutical preparation Mutaflor® by Alfred Nissle, after isolating it from the feces of a WWI soldier and characterizing its antagonistic effects against pathogenic *Enterobacterales* (Nissle, 1919). To date, Mutaflor® is a commercially available drug against intestinal diseases like Crohn’s disease and ulcerative colitis (Pharma-Zentrale GmbH, 2020).

While the microbial characteristics (Blum et al., 1995) and the genetic background (Grozdanov et al., 2004; Reister et al., 2014) of *Ec*N are well-studied and determined, the mechanism of probiotic activity remains elusive. The protective effects of *Ec*N are considered to be linked to several particular properties and fitness factors. *Ec*N secretes two bactericidal microcins, which have been shown to drive microbial competition (Sassone-Corsi et al., 2016; Vassiliadis et al., 2010). Biofilm formation as well as outcompeting pathogens regarding iron uptake (Deriu et al., 2013) are thought to be factors contributing to the probiotic properties of *Ec*N (Boudeau et al., 2003).

*Ec*N has shown high colonization success in gnotobiotic rats (Lorenz and Schulze, 1996) and piglets (Mandel et al., 1995) upon oral exposure, while also most conventionally kept mice and rats retain a long-term *Ec*N colonization of their guts (Schulze et al., 1992). In humans, however, only certain specific conditions allow stable, long-term colonization of the intestines. An extensive clinical study on healthy adults by Kurtz *et al.* (2018) suggests that a natural intestinal microbiome might interfere with *Ec*N preventing a long-term colonization and that the kinetics of *Ec*N in the human system are highly variable.

In addition to the importance for colonization, the communication of *Ec*N with epithelial cells has been shown to have numerous effects on the host’s immune system (Hafez et al., 2010; Sabharwal et al., 2016; Thomas and Versalovic, 2010). Well-studied immunomodulatory effects are the induction of the antimicrobial peptide human β-defensin-2 (HBD2) (Wehkamp et al., 2004) and the regulation of T-cell activation, expansion and apoptosis (Guzy et al., 2008; Sturm et al., 2005; Thomas and Versalovic, 2010). Directly interacting with the intestinal epithelial cells, *Ec*N is proposed to prevent a compromised epithelial barrier, which is considered a key mechanism in the development of intestinal diseases (Ukena et al., 2007). *Ec*N has been shown to inhibit “leaky gut” symptoms, which are associated with diseases like coeliac or Crohn’s disease, by enhancing the integrity of the mucosal surface of the intestinal epithelium (Souza et al., 2016; Ukena et al., 2007; Zyrek et al., 2007).

Although application of *Ec*N is usually safe, a few studies call for careful usage under specific conditions (Dziubańska-Kusibab et al., 2020; Gronbach et al., 2010; Günther et al., 2010). As a safety relevant aspect, not much is yet known about the phenotypic and genomic plasticity of *Ec*N under different growth conditions that may contribute to bacterial adaptation to different hosts or environments.

While mammals remain the most widely used model systems in research on infectious diseases and host-microbe interactions, alternative models might provide important benefits and are increasingly recognized as promising alternatives (Adiba et al., 2010; Lionakis, 2011). Among other factors, high costs, time-consuming maintenance, and the ethical concerns of infecting mammals with pathogens drive the substitution of insect models for mammals as host systems (Cutuli et al., 2019; Perini et al., 2019; Scully and Bidochka, 2006). Regarding microbial pathogenesis, insects and mammals show certain parallels in the structure of protective tissues (reviewed in Scully & Bidochka, 2006) and they share a highly conserved innate immune system with analogous signaling pathways (Heine and Lien, 2003; Hoffmann et al., 1999).

*Tribolium castaneum* is an established model organism and represents the benefits that insect model systems come with: *T. castaneum* has a short life cycle, high fecundity and can be inexpensively maintained, allowing experiments with large sample sizes (Sokoloff, 1977). It allows for RNAi studies using specific and efficient knockdowns of genes by larval or parental dsRNA injection (Bucher et al., 2002; Tomoyasu and Denell, 2004). As a member of the largest eukaryotic order, Coleoptera, *T. castaneum* makes a more suitable representative of other insects, since it is evolutionary more basal than Lepidoptera and Diptera (Misof et al., 2014). That makes the red flour beetle a commonly used insect model in various fields of research including development, evolution, immunity and host-pathogen interactions (Altincicek et al., 2008; Brown et al., 2009; Milutinović et al., 2013; Suzuki et al., 2008). Moreover, identification of various extra- and intracellular signaling pathways of the innate immune system, including many proteins with human homologs (Zou et al., 2007) allows for extensive and conclusive studies of *T. castaneum* immune system and its interactions with microbes. *T. castaneum* and its pathogens have become well-established models, in particular for immuno-ecological and evolutionary studies (e.g., Bérénos et al., 2011; Blaser & Schmid-Hempel, 2005; Ferro et al., 2019; Khan et al., 2017; Milutinović et al., 2013; Roth et al., 2009; Tate et al., 2017). Additionally, *T. castaneum* has been proposed as a screening system for potential drugs as well as pharmaceutical side effects (Bingsohn et al., 2016; Brandt et al., 2019). Grau et al. (2017) used *T. castaneum* for *in vivo* characterization of a probiotic *Enterococcus mundtii* isolate from *Ephestia kuehniella* larvae and suggested *T. castaneum* as an alternative model for the pre-screening of probiotics.

We here aimed to make use of these strengths of the *T. castaneum* system to further investigate the possible adaptation of *Ec*N to a new host and its environment. To better understand phenotypic and genomic plasticity of *Ec*N as mechanisms that can contribute to bacterial host adaptation, a serial passage experiment was performed, using multiple cycles of intestinal colonization in beetle larvae (Figure S1; Supplementary data). We observed a significant effect of eight serial passages on growth, motility and colony morphology of the passaged lines, suggesting that *Ec*N could adapt to novel environments. However, we did not observe any changes on the genomic level, suggesting that adaptation had happened on the phenotypic and/or epigenetic level.

Furthermore, we assessed the effects of *Ec*N treatment on the host system by monitoring life history traits (survival, pupation, eclosure) and measuring expression levels of antimicrobial peptides (AMPs) in *T. castaneum* larvae after oral exposure. Additionally, a putative protective effect against the beetle’s natural pathogen *Bacillus thuringiensis tenebrionis (Btt)* was analyzed by coinfecting larvae with *Ec*N and *Btt.* Even though we did not observe any effect on the host, changes in phenotypic attributes suggest that *Ec*N could phenotypically adapt to novel environments or unusual hosts, such as during the intestinal passage in *T. castaneum* larvae.

## Results

### Serial passages of *Ec*N had no effect on persistence in the host but changed phenotypic characteristics of *Ec*N

#### Persistence of EcN in T. castaneum did not increase after serial passages

A central aim of the serial passage experiment was to establish *Ec*N lines that are able to stably colonize the *T. castaneum* host. For the ancestral *Ec*N, preliminary experiments showed an exposure-time-dependent *Ec*N persistence of 4 to 7 days in *T. castaneum* larvae, upon oral exposure (Figure S2; supplementary data). Originating from one ancestral clone, 10 *Ec*N lines were passaged (L1 - 10) in the host (larvae-passaged), while 6 lines served as controls for the larval environment (flour-passaged) without being exposed to *T. castaneum.* We exposed 48 naïve larvae to each of the lines by keeping them separately in wells of 96-well plates for 48 h. We expected to select for persistence in the serial passage. We thus focused on measuring the persistence of the passaged lines compared to the ancestral strain after completing eight infection cycles. The persistence of the passaged lines was compared to the ancestral strain by testing individual larvae for the presence of *Ec*N.

To analyze the persistence of the serially passaged *Ec*N in the *T. castaneum* host, we performed an infection experiment that included all larvae-passaged *Ec*N lines after 8 passages, as well as the ancestral strain. After 48 h on an *Ec*N diet, we transferred all larvae to flour diet without *Ec*N and sampled for the following 7 days, a subset of 20 larvae per replicate each day (Transparent methods; supplementary data). We did not detect a significant difference in the ability to persist in the beetle larvae between the ancestral strain and any of the passaged lines (Figure 1). One day after exposure, the passaged *Ec*N lines varied in persistence in the beetle larvae with an average persistence rate of 46 %, while 75 % of the tested larvae treated with the ancestral strain harbored *Ec*N. Over time, the rate of larvae harboring *Ec*N decreased, and after 7 days, the mean persistence rate was 5 %. In two lines (L9 and L10) *Ec*N was not found anymore.

**Figure 1.**
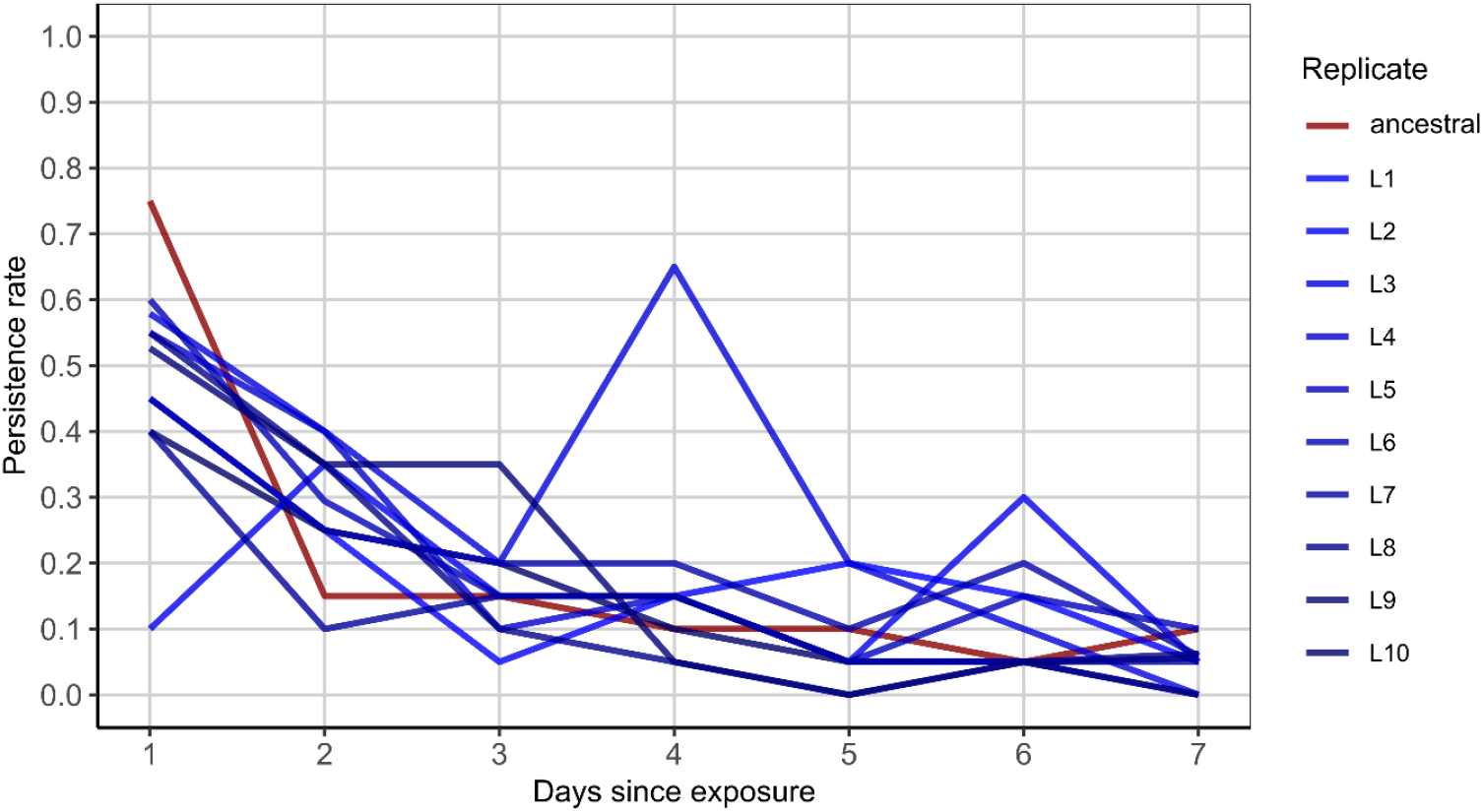
Persistence of *Ec*N in *T. castaneum* larvae after serial passage. Proportion of larvae harboring *Ec*N after 48 h of exposure. Ten passaged *Ec*N lines (L1-10) and the ancestral strain were monitored. 20 larvae per replicate and day were tested for bacterial presence or absence (n = 1584).

#### Host and flour-passaged lines showed increased growth rate compared to ancestral EcN

To characterize putative trade-offs resulting from adaptation to the new environment upon serial passage, we investigated potential effects on bacterial growth parameters. The absorbance of liquid cultures inoculated with a standardized number of cells was monitored for 24 h (Figure 2). Six of the larvae- and flour-passaged replicate lines as well as 6 ancestral clones (pseudo-replicates) were measured every 15 min.

**Figure 2:**
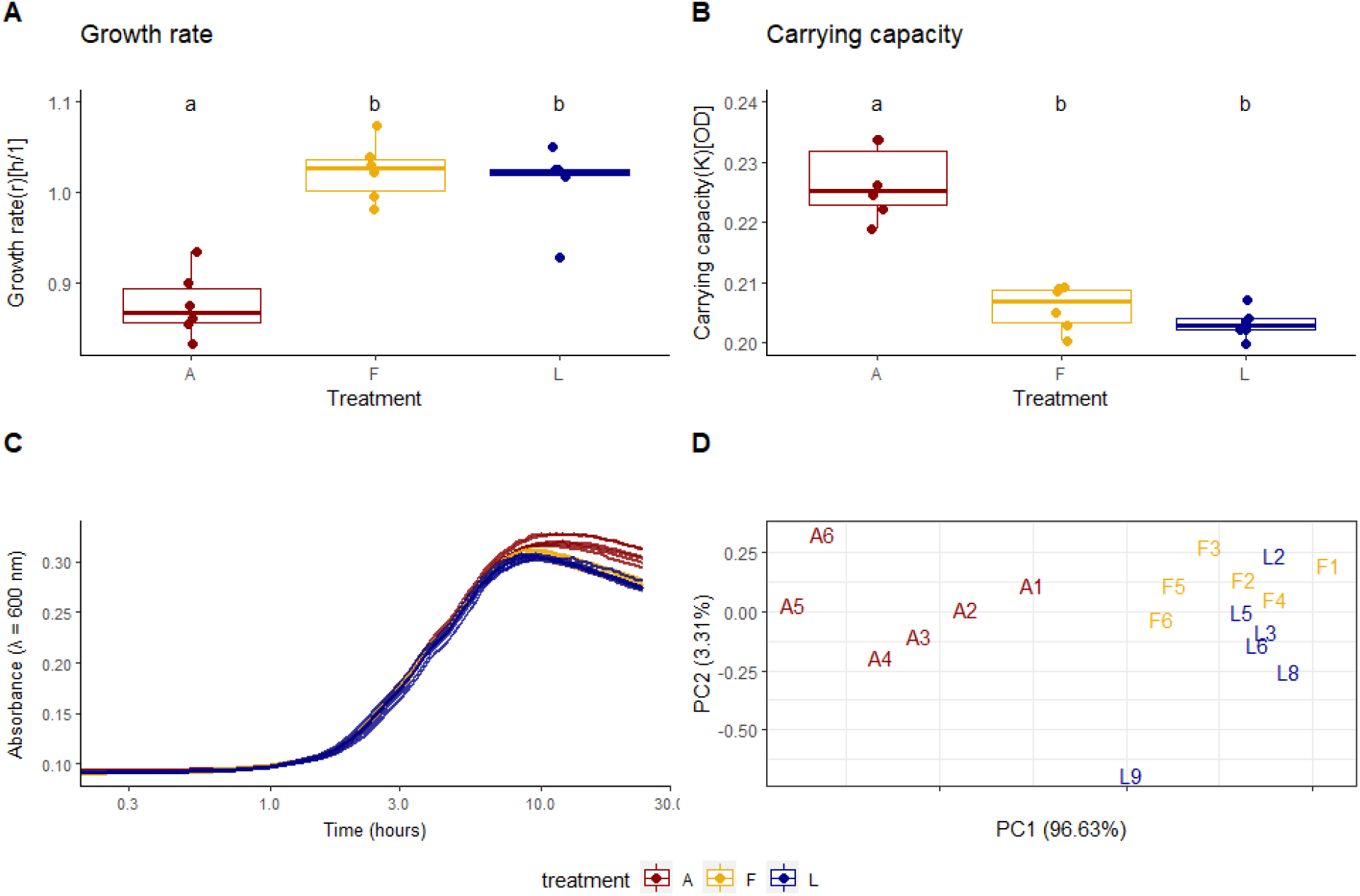
*Ec*N growth curves and attributes. A) Differences in the growth rates between the passaged lines and the ancestral strain (Kruskal-Wallis *X*^*2*^ = 10.889, Df = 2, p = 0.00432). B) Differences in the carrying capacities (Kruskal-Wallis *X*^*2*^ = 12.316, Df = 2, p-value = 0.00211). C) Optical density (absorbance) of bacterial liquid cultures at λ = 600 nm. D) Principal component analysis of growth rate, carrying capacity, generation time and the area under the curve. Six replicates per treatment were analyzed: Larvae-passaged (L2, L3, L5, L6, L8, L9), flour-passaged (F1-6), ancestral (A1-6, pseudo-replicates). Measurements were taken every 15 min for 24 h at 30 °C. A - ancestral strain, F - flour-passaged bacteria, L - larvae-passaged bacteria. Statistical differences between the treatments are marked with letters a and b.

The curves of all lines follow a similar, logarithmic growth (Figure 2C). The maximum or intrinsic growth rate (r) turned out to be significantly higher in the evolved lines (Figure 2A), while the ancestral lines appeared to grow to a higher density than the passaged lines. Analysis of the growth capacities (K) confirmed a significantly higher maximum density in the ancestral lines (Figure 2B).

We performed a principal component analysis (PCA) of r and K, as well as generation time and the area under the curve to visualize similarities and differences in these attributes between the individual lines (Figure 2D). The area under the curve is a conclusive parameter because it integrates the contributions of the growth rate, carrying capacity, and the initial population size, into a single value. The PCA revealed that the growth parameters of the ancestral lines appear distinct from the evolved ones, which cluster together. The passaged line L9, however, showed distinctive growth characteristics dissimilar to all other lines (Figure 2D).

#### Larvae- and flour-passaged EcN lines were more motile than the ancestral strain

Differences in the motility of bacterial lines can, similar to growth, indicate trade-offs linked to a putative adaptation to the novel larval environment. Therefore, we assessed the swarming distance on soft agar plates of 3 technical replicates of 6 of the larvae-passaged (L) and flour-passaged lines (F) as well as 6 pseudo-replicates of the ancestral strain (A) after 6 h (Figure 3A) and 9 h (Figure 3B). Out of the 10 larvae-passaged lines we chose 6 that showed a trend of elevated persistence.

**Figure 3:**
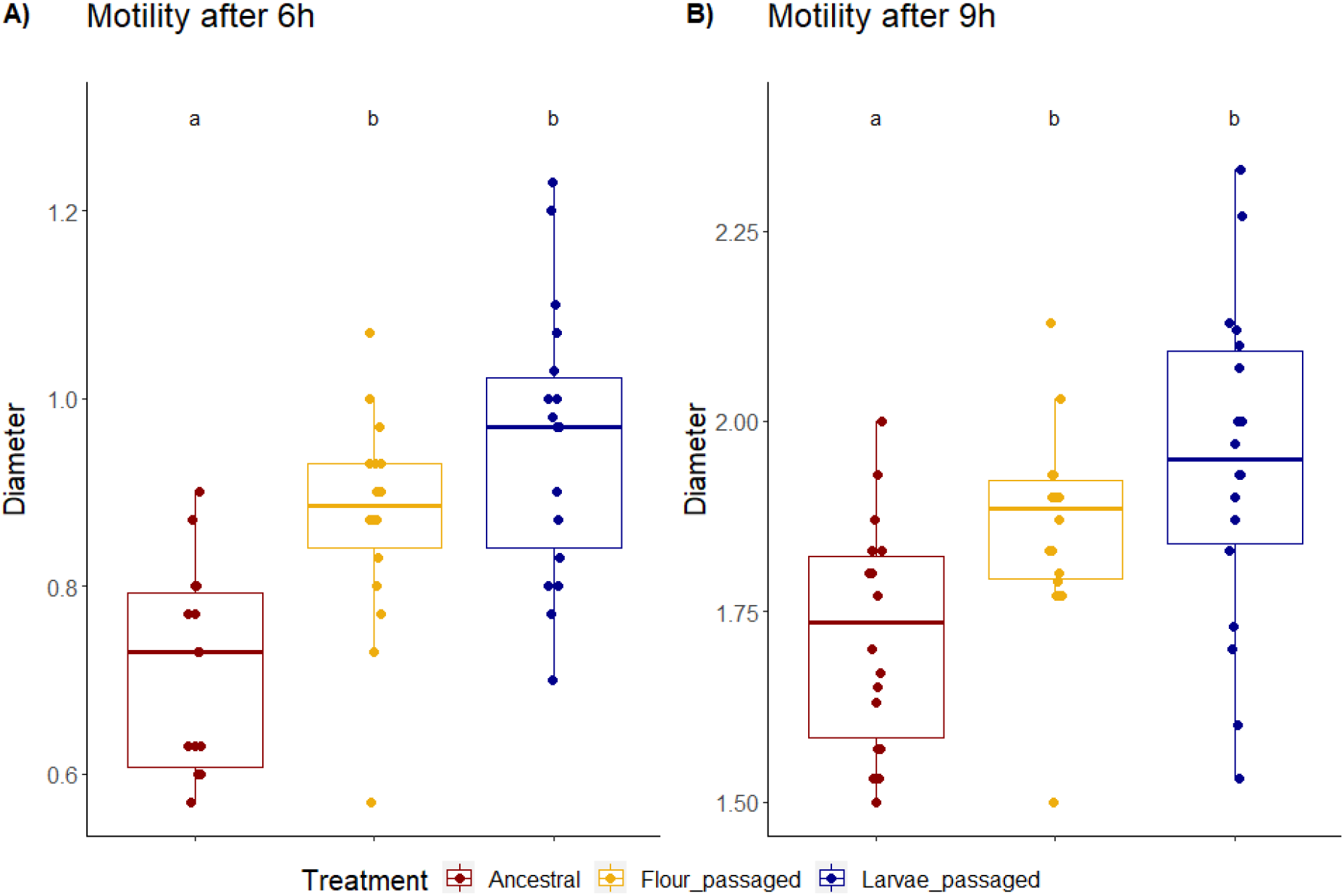
*Ec*N motility. Differences in swarming ability between the evolved lines and the ancestral strain. The radius of the swarmed areas on 0.3 % agar plates was measured after 6 h (3 A) and 9 h (3 B) post inoculation. Triplicates of six replicates per treatment were analyzed: Larvae-passaged (L2, L3, L5, L6, L8, L9), flour-passaged (F1-6), ancestral (A1-6, pseudo-replicates). 3A is showing motility per treatment after 6 h (anova, Df=2, F=17.4, p <0.001). 3B is showing flagella motility after 9h per treatment (anova, Df=2, F=8.06, p <0.001). Statistical differences between the treatments are marked with letters a and b.

After 6 h at 30 °C the larvae-passaged lines, as well as the flour lines showed significantly higher motility than the ancestral strain (average swarming distance: L passaged lines: 4.76 mm, F passaged lines: 4.36 mm, A strain: 3.58 mm). This difference was still observed after 9 h of incubation. No significant difference between the passaged lines was detected but the larvae-passaged lines tended to be more motile than the flour-passaged lines at both time points. Despite the elevated motility of the passaged lines, the line L9 showed particularly low average swarming distances of 3.78 mm after 6h and 8.06 mm after 9 h (Figure S3; Supplementary data)

#### Colony morphology of one replicate line changed after passages through the host

To investigate possible differences in colony morphology we grew three colonies of the following replicate lines on dye supplemented agar plates: Larvae-passaged (L2, L3, L5, L6, L8, L9), flour-passaged (F1-6), ancestral (A1-6, pseudo-replicates). The synthesis of cellulose and curli fimbriae, components of the extracellular matrix were visualized using the dyes Calcofluor White (under UV) and Congo Red, respectively (Cimdins and Simm, 2017; Zogaj et al., 2001). While *E. coli* K-12 colonies generally produce more curli fimbriae than cellulose at 30 °C, thus appearing as brown and smooth colonies, *Ec*N displays a pdar (pink, dry and rough) colony morphotype (Cimdins and Simm, 2017; Grozdanov et al., 2002). One of the passaged lines, L9, appeared to display a slightly stronger fluorescence upon Calcofluor White staining, in addition to a more wrinkled colony outline, which may indicate increased cellulose expression (Figure 4).

**Figure 4:**
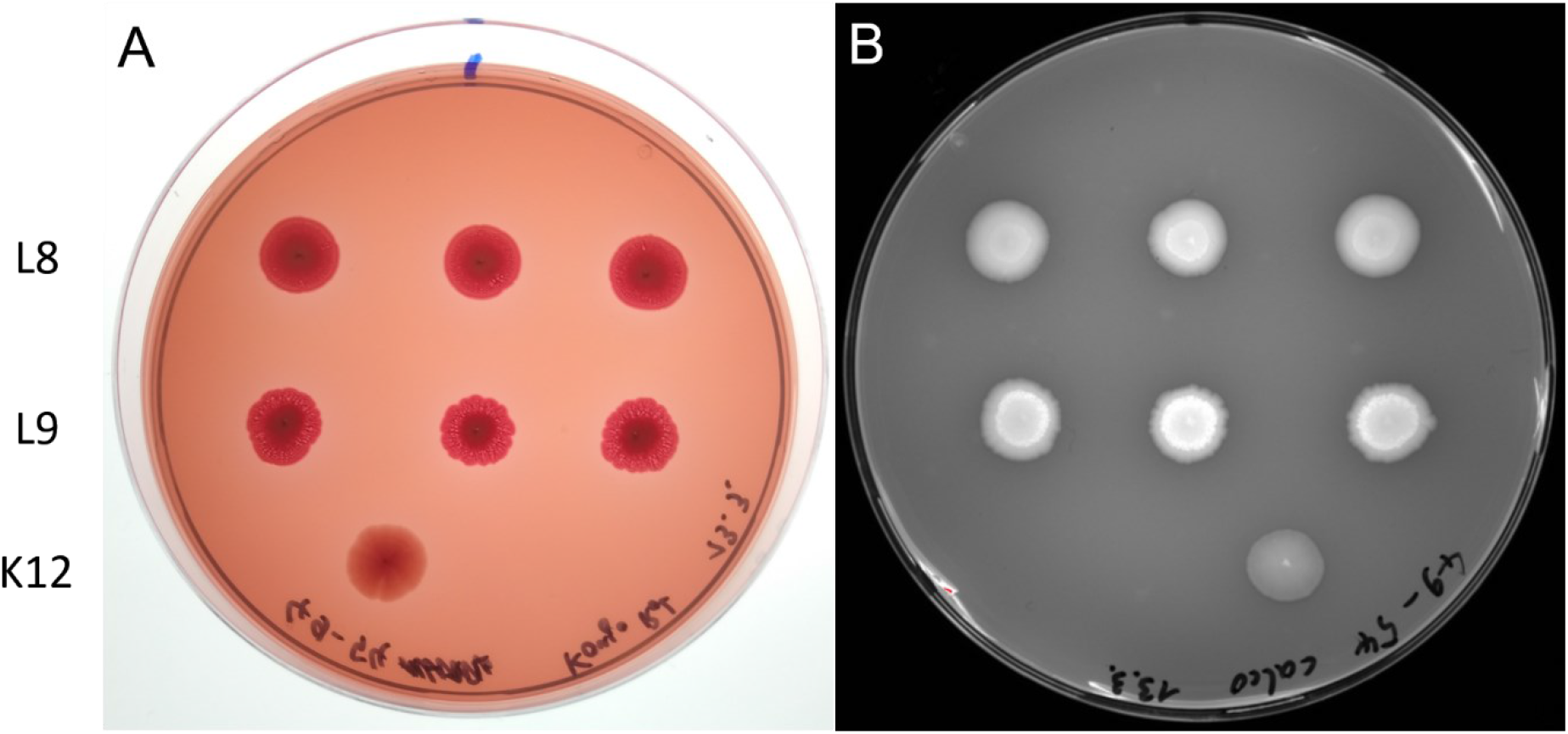
*Ec*N L9 colony morphology. *Ec*N colonies grown on Congo Red (left) and Calcofluor White plates (right). Triplicates of six replicates per treatment were tested: Larvae-passaged (L2, L3, L5, L6, L8, L9), flour-passaged (F1-6), ancestral (A1-6, pseudo-replicates). Triplicates of the lines L8 (top row) and L9 (middle row) are shown. *E. coli* K-12 MG1655 served as a control (bottom). The plates were grown for 96 h at 30 °C. Morphology of the line L8 showed typical *Ec*N morphology.

### Genome of *Ec*N remained unchanged after serial passage

The morphology and the strongly diminished growth and motility of the host-passaged line L9 suggested that a mutation might have occurred and defined these properties. Therefore, we analyzed the genomic variabilities of the draft genome sequences of this and two other lines, which showed elevated motility (L2, F4). Both variant detection and pan-genome analysis of these isolates showed the absence of genome-level differences compared to the ancestral strain.

### *Ec*N did not influence host mortality or gene expression

#### EcN did not provide a survival benefit to the host upon pathogen exposure

*Ec*N shows probiotic activity in humans, and we were thus interested in its putative beneficial effects on the survival and development of *T. castaneum* larvae upon *Btt* infection. Beetle larvae were first orally exposed to *Ec*N (‘pretreatment’), followed by oral exposure to *Btt* 3 days later (‘treatment’). *Ec*N for the pretreatment were derived from the ancestral strain or either of three of the larvae-passaged *Ec*N lines. For this, lines L2, L3, and L6 were selected, because they showed a tendency for above-average persistence (Figure 5). As additional controls, the commensal *E. coli* K-12 strain MG1655 (K12) served as a general control for bacterial pretreatment, and unexposed PBS controls were included for both pretreatment and treatment.

**Figure 5:**
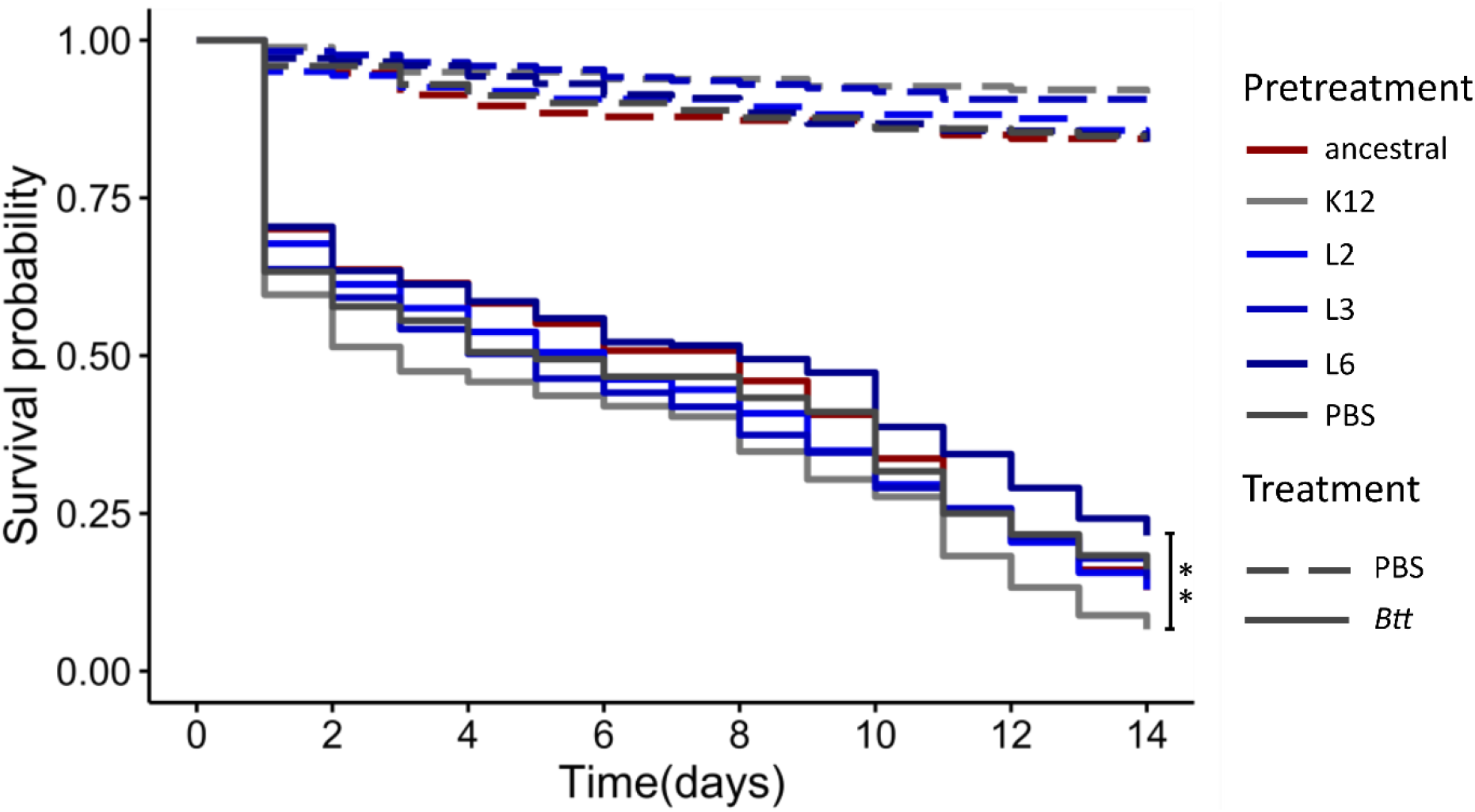
*T. castaneum* survival upon *Ec*N pretreatment. Survival of 14 day-old beetle larvae exposed to *Ec*N-containing flour diet for 72 h (5.3 × 10^10^ cells / g flour), before transfer to *Btt*-containing flour (3.3 × 10^10^ spores / g flour). Three passaged *Ec*N strains (L2, L3, L6) as well as the ancestral strain were used for pretreatment. PBS and K-12 strain MG1655 served as a negative control for pretreatment and treatment. The larvae were individualized in 96-well plates (n = 2128). ** p = 0.0046. Full lines show survival of larvae challenged with *Btt* while dashed lines are PBS control.

The *Btt* infection strongly reduced larval survival (Figure 5), whereas the *Ec*N pretreatment did not have any strong effect on larval survival of the *Btt* infection. Only larvae pretreated with one of the host-passaged *Ec*N lines (line 6) showed a slightly increased survival, while *E. coli* K-12 pretreatment slightly reduced survival, an effect that was significant only in direct comparison of these groups (*χ2* = 16.91, Df = 5, p=0.0046) but not when compared to the PBS control.

In addition to assessing the effects of *Ec*N exposure on survival, we monitored the pupation and eclosure rates of the larvae for 4 weeks but did not observe any significant differences (Figure S4, S5; Supplementary data).

#### Expression of immune-related genes did not change in Tribolium castaneum upon oral uptake of EcN

We assessed the expression levels of several immune genes by RT-qPCR to characterize a possible differential immune reaction of *T. castaneum* to the exposure to ancestral *Ec*N. The selected genes code for the AMPs Attacin2, Cecropin2, Defensin2, and Defensin3 (Yokoi et al., 2012), as well as for an Osiris16-like protein, which was found to be involved in oral immune priming with *Btt* (Greenwood et al., 2017). The expression patterns were measured in larvae, after oral exposure to ancestral *Ec*N or *E. coli* K-12 (as a control) for 72 h (6 replicates of 5 pooled larvae per treatment). We did not find any statistically significant differences in gene expression to PBS-exposed control larvae (Figure 6).

**Figure 6:**
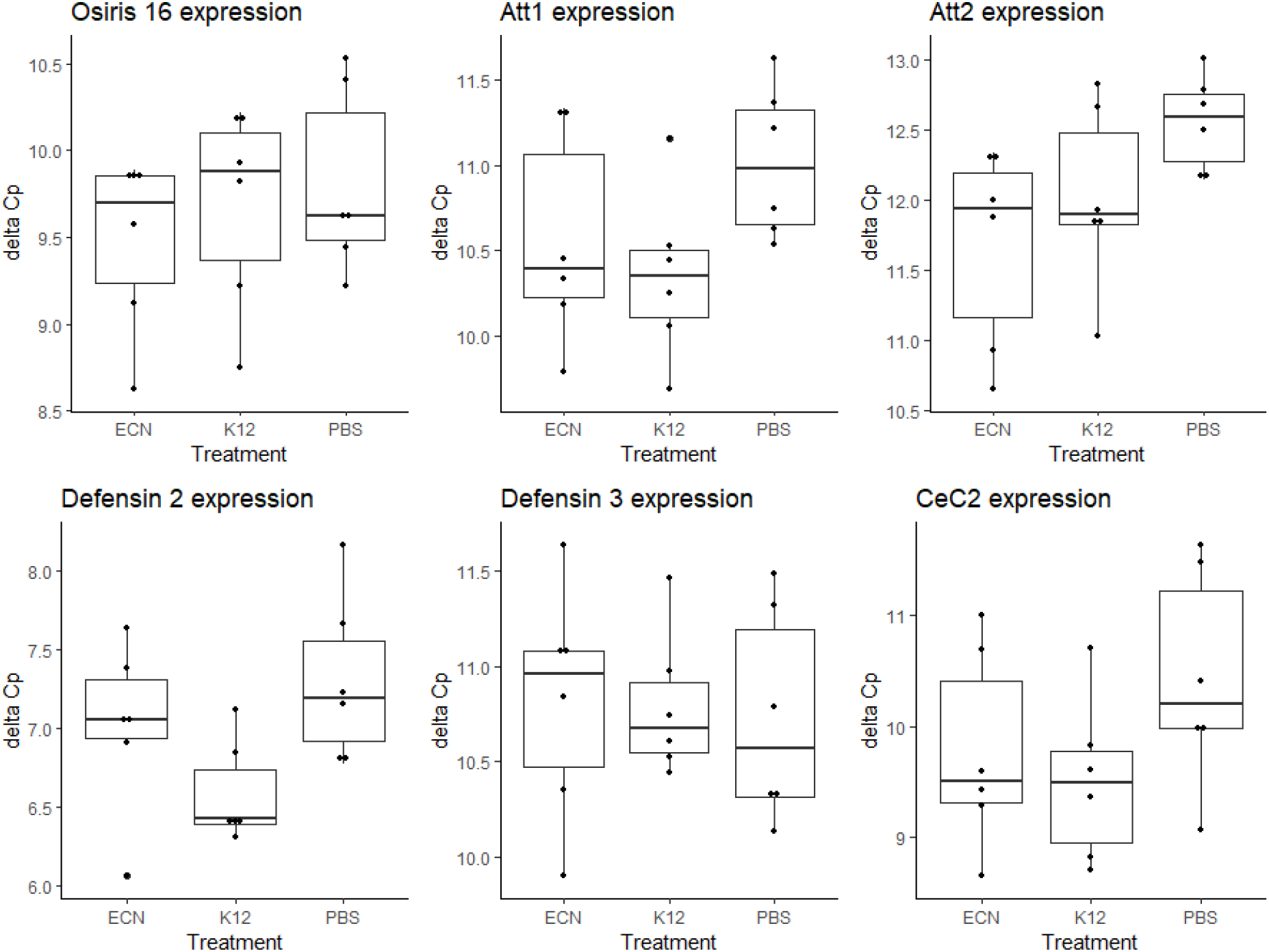
Differential expression of AMPs and Osiris16. Expression patterns assessed by RT-qPCR on RNA extracted from 6 replicates of 5 pooled larvae/treatment. *Tribolium castaneum* larvae that were orally exposed to *Ec*N, *E. coli* K-12 MG1655 and PBS for 72 h. The genes coding for the AMPs Attacin2 (Att2), Cecropin2 (Cec2), Defensin2 (Def2) and Defensin3 (Def3) as well as for an Osiris16-like protein were analyzed. PBS treatment served as a negative control. ΔCp values were calculated using the expression of the housekeeping genes ribosomal protein L13a (Rpl13a) and ribosomal protein 49 (Rp49).

## Discussion

Serial passage experiments are important tools to understand the evolution of bacteria in new environments, in particular new hosts, and to adapt bacteria to new experimental host systems. We performed a serial passage experiment of the human probiotic *Ec*N in an invertebrate host, larvae of the red flour beetle *T. castaneum*, as well as in their environment and food source, i.e., flour. We observed clear changes in key phenotypic parameters of the bacteria, which surprisingly did not lead to increased persistence of the bacteria in this novel host, nor to any genetic changes.

By extracting *Ec*N after an increasingly long duration in the larval system, selection for persistence was expected, but the selection pressure exerted by the novel environment might not have been sufficient. Additionally, the selection pressure exerted by desiccation in the flour might outweigh the effects of the serial passage in the host.

Both larvae and flour-passaged lines showed increased growth rates in liquid culture, but a decreased carrying capacity relative to the ancestral strain (Figure 2). Somerville et al. (2002) also found significantly elevated growth rates in *Staphylococcus aureus* upon serial passage *in vitro* and suggested that the difference was based on the more efficient utilization of available nutrients upon serial passage. Likewise, the nutrient-scarce environment of the flour and the larval gut during passage might have selected for efficient nutrient uptake, leading to increased growth, in our experiment. The reduced carrying capacity could be caused by a trade-off for the increased growth rate.

The assessment of motility upon passage showed a significantly higher swimming capability on soft agar for the evolved lines compared to the ancestral strain (Figure 3). Taxis of *E. coli* bases on the expression of flagella and is thought to be very costly, leading to infrequent expression (Soutourina and Bertin, 2003). The biosynthesis of flagella has been shown to be strongly regulated by environmental factors including pH, salinity, temperature, and presence of D-glucose (Li et al., 1993; Soutourina et al., 2002). Moreover, Landini & Zehnder (2002) demonstrated that the flagellar motility of *E. coli* is induced by oxygen-limited conditions. Thus, the difference in motility between the passaged lines and the ancestral strain might have resulted from the extreme environmental conditions during the passage, which could have promoted the expression of flagella. Apart from motility, flagella play a major role in the adhesion of *Ec*N to host cells and biofilm formation (Kleta et al., 2014).

The larvae-passaged line L9 appeared to display a slightly stronger fluorescence upon Calcofluor White staining, in addition to a more wrinkled colony outline (Figure 4). An increased fluorescence on Calcofluor White suggests a higher cellulose concentration in the extracellular matrix, which is linked to the wrinkled colony phenotype (Zogaj et al., 2001). Since L9 is one of the lines that were passaged through the host, it is conceivable, that the novel phenotype arose in an adaptive process to the larval environment. However, the low persistence of L9 indicates that this phenotype is not associated with colonization success (Figure 1). Nevertheless, the peculiar features of L9 are a proof-of-principle that serial passage of *Ec*N can generate differential phenotypic characteristics. The morphology and the strongly diminished growth and motility of the larvae-passaged line L9 suggested that a mutation might have occurred and defined these properties. However, draft whole genome sequences of this and two other lines, which showed elevated motility (L2, F4) did not reveal any genetic difference to the parental strain, suggesting possible epigenetic variations.

Oral previous exposure of *T. castaneum* larvae to serially passaged or ancestral *Ec*N, prior to exposure to the entomopathogen *B. thuringiensis tenebrionis (Btt)*, did not significantly improve host survival (Figure 5). Only one of the larvae-passaged lines (L6) appears to induce a slightly elevated survival rate, compared to pretreatment with *E. coli* K-12 strain MG1655. In conclusion, *Ec*N does not seem to act as a probiotic in *T. castaneum* that protects against pathogens, at least not against *Btt*.

The antagonistic activity of *Ec*N against pathogens might need specific requirements and an unaccustomed microbiome, as well as suboptimal environmental conditions in the beetle gut might have impaired its probiotic effect in the novel host. For example, iron homeostasis has been shown to play a major role in infectious diseases and successfully competing for iron is thought to be a central mechanism of the antagonistic effect of *Ec*N against pathogens (Deriu et al., 2013; Weiss, 2013). Furthermore, Sassone-Corsi *et al.* (2016) demonstrated that elevated concentrations of environmental iron inhibit the microcin production of *Ec*N. Moreover, the bactericidal activity of *Ec*N’s siderophore microcins M and H47 has been shown to be restricted to members of the *Enterobacteriaceae* family with no activity against gram-positive bacteria, including *Bacillus* (Vassiliadis et al., 2010).

Another probiotic effect of *Ec*N could be the enhancement of the integrity of the gut epithelium. In mammals, *Ec*N induces the upregulation of the structural protein Zonula occludens-1 (ZO-1), which binds to both, actin filaments and the tight junction protein occludin, structurally linking tight junctions to the cytoskeleton (Ukena et al., 2007; Wittchen et al., 1999; Zyrek et al., 2007). The insect homolog of ZO-1 is the *Drosophila* discs-large tumor suppressor protein (Dgl), which has been shown to have a similar function to that of ZO-1 in septate junctions, the invertebrate homolog of tight junctions (Harden et al., 2016; Willott et al., 1993). ZO-1 and Dgl, however, show distinct differences in the secondary and tertiary protein structure and the regulation of expression might differ considerably despite the homology in function (Willott et al., 1993). These dissimilarities could adversely affect or completely impair the stabilizing effect of *Ec*N on the epithelial permeability of *T. castaneum*.

The absence of any effects on survival, pupation and eclosure of beetles after *Ec*N exposure as well as expression levels of several tested immune genes (mostly AMPs; Figure 6) suggest that there was no impact on the immune response and development of *T. castaneum.* Yokoi et al. (2012) investigated the expression of AMPs upon exposure to multiple microorganisms including *E. coli* strain DH5α and found all of the AMPs that were tested in the present study upregulated. However, the effect was observed in pupae and the microorganisms used in their study were directly injected, instead of oral exposure (Yokoi et al., 2012). However, similar immune genes as tested here (Osiris 16) were differentially regulated upon oral infection or priming of *T. castaneum* with the entomopathogen *Btt* (Behrens et al., 2014; Greenwood et al., 2017), pointing towards less immunogenic effects of *Ec*N upon oral exposure.

In conclusion, this exploratory study did not show any increased colonization success nor protective effect of *Ec*N upon 8 serial passages through *T. castaneum* larvae. However, we observed phenotypic changes in growth attributes and motility in lines that were passaged through the host as well as those passaged through flour only. Moreover, the peculiar characteristics of one of the lines (L9) show that serial passage of *Ec*N can generate differential phenotypes, even in the absence of changes on the genomic level. A longer duration of the serial passage experiment might be necessary to lead to genomic changes and yield further valuable insight into the adaptation of *Ec*N to a novel host environment.

## Limitations of the study

One of the main goals of this project was the establishment of a novel model system for studying the human probiotic *Ec*N. An invertebrate system could facilitate research on this widely used probiotic. However, we were not able to increase persistence of the passaged *Ec*N lines. A probable reason is an insufficient number of passages, given the reported low intrinsic mutation rates of *E. coli*: 4.1 × 10^−4^ for strain REL606; (Wielgoss et al., 2011) to 1.0 × 10^−3^ mutations per genome per generation for K-12 strain MG1655 (Lee et al., 2012). Estimating the number of generations that the bacterial lines went through in the performed serial passage experiment is rather difficult since it is not apparent if and how fast *Ec*N replicates within the host or the flour. Jerome et al. (2011) showed that adaptation of a human intestinal bacterium to a novel mouse model host is possible within as few as 3 passages but suggest that a fundamental ability to colonize a broad host spectrum and high standing genetic diversity may be crucial to long term persistence. We started our serial passages with a single clone eliminating standing genetic diversity, which selection could have acted on.

The sensitivity and precision of the plating method that was used to measure persistence after serial passage are not determined and might have been insufficient for a conclusive analysis. Due to a limited number of exposed larvae, the persistence was only observed for 7 days after exposure, which did not allow for assessing differences in long-term colonization success.

Despite the strong phenotypic changes that line L9 shows after serial passage through the host, we do not know how stable these changes are, since they were not tested after culturing without selection in the host or flour. Epigenetic mechanisms such as inheritance of DNA methylation patterns are recognized in bacterial biology (Casadesús and Low, 2006). Considering we did not observe any genomic differences, we are suggesting that observed phenotypic differences could arise from epigenetic processes, but further analysis should be done to confirm this.

Differences between vertebrate and invertebrate physiology are limitations that could prevent the attempted establishment of invertebrate model systems for vertebrate probiotics. Despite parallels between mammals and insects in how they deal with microorganisms, the immunological and physiological differences between the intestinal environments of *T. castaneum* and humans might prevent successful adaptation of *Ec*N to an invertebrate gut. One of the mechanisms important for *Ec*N colonization is binding to the host’s gastrointestinal mucus with the flagellum as the major adhesin (Troge et al., 2012). Only recently, Dias et al. (2018) showed that several insects produce mucous substances, including the mealworm beetle *Tenebrio molitor*, a close relative of *T. castaneum*. However, mammals and insects differ strongly in chemical and enzymatic composition of the gastrointestinal mucus layer (Dias et al., 2018; Juge, 2012). Moreover, the differing body temperature of the host species might be another important factor preventing adhesion and growth in the host gut (Gill et al., 2019; Moghadam et al., 2018).

## Resource of availability

Lead contact

J. Kurtz ( is the lead contact for this paper.

Materials

All passaged *Ec*N lines are available upon request from J. Kurtz.

## Data and code availability

All data and R scripts used to run statistical analyses and produce the figures discussed in this paper are deposited in the following GitHub repository: https://github.com/Bio-nic/serial_passage_Nissle

The genome sequencing data will be uploaded to the NCBI BioProject database under the following accession: PRJNA684051

## Methods

All methods can be found in the Transparent methods file submitted along with the manuscript.

## Acknowledgments

We would like to thank Kathrin Brüggemann for performing RNA extractions and qPCR and Olena Mantel for her support and work on phenotypic assays. This work was carried out within the DFG Research Training Group 2220 “Evolutionary Processes in Adaptation and Disease” at the University of Münster (281125614/GRK 2220). Haleluya Wami was supported by a scholarship by Pharma-Zentrale GmbH.

## Authors contributions

Conceptualization, N.C.H.S., A.K., J.K.; Investigation, N.C.H.S., A.K.; Data analysis: A.K., N.C.H.S., H.W.; Writing the original draft: N.C.H.S, A.K.; Editing and writing, N.C.H.S., A.K., H.W, J.K., U.D.; Funding acquisition: J.K and U.D.

## References

Adiba, S., Nizak, C., van Baalen, M., Denamur, E., Depaulis, F., 2010. From grazing resistance to pathogenesis: The coincidental evolution of virulence factors. PLoS One. https://doi.org/10.1371/journal.pone.0011882

Altincicek, B., Knorr, E., Vilcinskas, A., 2008. Beetle immunity: Identification of immune-inducible genes from the model insect Tribolium castaneum. Dev. Comp. Immunol. 32, 585–595. https://doi.org/10.1016/j.dci.2007.09.005

Behrens, S., Peuß, R., Milutinović, B., Eggert, H., Esser, D., Rosenstiel, P., Schulenburg, H., Bornberg-Bauer, E., Kurtz, J., 2014. Infection routes matter in population-specific responses of the red flour beetle to the entomopathogen Bacillus thuringiensis. BMC Genomics 15, 445. https://doi.org/10.1186/1471-2164-15-445

Bérénos, C., Wegner, K.M., Schmid-Hempel, P., 2011. Antagonistic coevolution with parasites maintains host genetic diversity: An experimental test, in: Proceedings of the Royal Society B: Biological Sciences. Royal Society, pp. 218–224. https://doi.org/10.1098/rspb.2010.1211

Bingsohn, L., Knorr, E., Vilcinskas, A., 2016. The model beetle Tribolium castaneum can be used as an early warning system for transgenerational epigenetic side effects caused by pharmaceuticals. Comp. Biochem. Physiol. Part - C Toxicol. Pharmacol. 185–186, 57–64. https://doi.org/10.1016/j.cbpc.2016.03.002

Blaser, M., Schmid-Hempel, P., 2005. Determinants of virulence for the parasite Nosema whitei in its host Tribolium castaneum. J. Invertebr. Pathol. 89, 251–257. https://doi.org/10.1016/j.jip.2005.04.004

Blum, G., Hacker, J., Marre, R., 1995. Properties of Escherichia coli strains of serotype O6. Infection 23, 234–236. https://doi.org/10.1007/BF01781204

Borges, S., Silva, J., Teixeira, P., 2014. The role of lactobacilli and probiotics in maintaining vaginal health. Arch. Gynecol. Obstet. 289, 479–489. https://doi.org/10.1007/s00404-013-3064-9

Boudeau, J., Glasser, A.-L., Julien, S., Colombel, J.-F., Darfeuille-Michaud, A., 2003. Inhibitory effect of probiotic Escherichia coli strain Nissle 1917 on adhesion to and invasion of intestinal epithelial cells by adherent-invasive E. coli strains isolated from patients with Crohn’s disease. Aliment. Pharmacol. Ther. 18, 45–56. https://doi.org/10.1046/j.1365-2036.2003.01638.x

Brandt, A., Joop, G., Vilcinskas, A., 2019. *Tribolium castaneum* as a whole‐animal screening system for the detection and characterization of neuroprotective substances. Arch. Insect Biochem. Physiol. 100, e21532. https://doi.org/10.1002/arch.21532

Brown, S.J., Shippy, T.D., Miller, S., Bolognesi, R., Beeman, R.W., Lorenzen, M.D., Bucher, G., Wimmer, E.A., Klingler, M., 2009. The red flour beetle, Tribolium castaneum (Coleoptera): A model for studies of development and pest biology. Cold Spring Harb. Protoc. 4, pdb.emo126. https://doi.org/10.1101/pdb.emo126

Bucher, G., Scholten, J., Klingler, M., 2002. Parental RNAi in Tribolium (Coleoptera). Curr. Biol.

Casadesús, J., Low, D., 2006. Epigenetic Gene Regulation in the Bacterial World. Microbiol. Mol. Biol. Rev. 70, 830–856. https://doi.org/10.1128/mmbr.00016-06

Cimdins, A., Simm, R., 2017. Semiquantitative analysis of the red, dry, and rough colony morphology of salmonella enterica serovar typhimurium and Escherichia coli using congo red, in: Methods in Molecular Biology. Humana Press Inc., pp. 225–241. https://doi.org/10.1007/978-1-4939-7240-1_18

Cutuli, M.A., Petronio, G., Vergalito, F., Magnifico, I., Pietrangelo, L., Venditti, N., Di Marco, R., 2019. Galleria mellonella as a consolidated in vivo model hosts: New developments in antibacterial strategies and novel drug testing. Virulence. https://doi.org/10.1080/21505594.2019.1621649

Deriu, E., Liu, J.Z., Pezeshki, M., Edwards, R.A., Ochoa, R.J., Contreras, H., Libby, S.J., Fang, F.C., Raffatellu, M., 2013. Probiotic bacteria reduce salmonella typhimurium intestinal colonization by competing for iron. Cell Host Microbe 14, 26–37. https://doi.org/10.1016/j.chom.2013.06.007

Dias, R.O., Cardoso, C., Pimentel, A.C., Damasceno, T.F., Ferreira, C., Terra, W.R., 2018. The roles of mucus-forming mucins, peritrophins and peritrophins with mucin domains in the insect midgut. Insect Mol. Biol. 27, 46–60. https://doi.org/10.1111/imb.12340

Dziubańska-Kusibab, P.J., Berger, H., Battistini, F., Bouwman, B.A.M., Iftekhar, A., Katainen, R., Cajuso, T., Crosetto, N., Orozco, M., Aaltonen, L.A., Meyer, T.F., 2020. Colibactin DNA-damage signature indicates mutational impact in colorectal cancer. Nat. Med. 26, 1063–1069. https://doi.org/10.1038/s41591-020-0908-2

Faubion, W.A., Sandborn, W.J., 2000. Probiotic therapy with E. coli for ulcerative colitis: take the good with the bad. Gastroenterology 118, 630–631. https://doi.org/10.1016/S0016-5085(00)70272-1

Ferro, K., Peuß, R., Yang, W., Rosenstiel, P., Schulenburg, H., Kurtz, J., 2019. Experimental evolution of immunological specificity. Proc. Natl. Acad. Sci. U. S. A. 116, 20598–20604. https://doi.org/10.1073/pnas.1904828116

Gill, J., Lau, K., Lee, T., Lucas, H., 2019. Escherichia coli Nissle 1917 forms biofilm and outgrows Escherichia coli K12 in a temperature-dependent manner, Undergraduate Journal of Experimental Microbiology and Immunology (UJEMI).

Grau, T., Vilcinskas, A., Joop, G., 2017. Probiotic Enterococcus mundtii isolate protects the model insect Tribolium castaneum against Bacillus thuringiensis. Front. Microbiol. 8. https://doi.org/10.3389/fmicb.2017.01261

Greenwood, J.M., Milutinović, B., Peuß, R., Behrens, S., Esser, D., Rosenstiel, P., Schulenburg, H., Kurtz, J., 2017. Oral immune priming with Bacillus thuringiensis induces a shift in the gene expression of Tribolium castaneum larvae. BMC Genomics 18, 1–14. https://doi.org/10.1186/s12864-017-3705-7

Gronbach, K., Eberle, U., Müller, M., Ölschläger, T.A., Dobrindt, U., Leithäuser, F., Niess, J.H., Döring, G., Reimann, J., Autenrieth, I.B., Frick, J.S., 2010. Safety of probiotic Escherichia coli strain Nissle 1917 depends on intestinal microbiota and adaptive immunity of the host. Infect. Immun. 78, 3036–3046. https://doi.org/10.1128/IAI.00218-10

Grozdanov, L., Raasch, C., Schulze, J., Sonnenborn, U., Gottschalk, G., Hacker, J., Dobrindt, U., 2004. Analysis of the genome structure of the nonpathogenic probiotic Escherichia coli strain Nissle 1917. J. Bacteriol. 186, 5432–5441. https://doi.org/10.1128/JB.186.16.5432-5441.2004

Grozdanov, L., Zähringer, U., Blum-Oehler, G., Brade, L., Henne, A., Knirel, Y.A., Schombel, U., Schulze, J., Sonnenborn, U., Gottschalk, G., Hacker, J., Rietschel, E.T., Dobrindt, U., 2002. A single nucleotide exchange in the wzy gene is responsible for the semirough O6 lipopolysaccharide phenotype and serum sensitivity of Escherichia coli strain Nissle 1917. J. Bacteriol. 184, 5912–5925. https://doi.org/10.1128/JB.184.21.5912-5925.2002

Günther, K., Straube, E., Pfister, W., Günther, A., Hübler, A., 2010. Severe Sepsis After Probiotic Treatment With Escherichia coli Nissle 1917. Pediatr. Infect. Dis. J. 29, 188–189. https://doi.org/10.1097/INF.0b013e3181c36eb9

Guzy, C., Paclik, D., Schirbel, A., Sonnenborn, U., Wiedenmann, B., Sturm, A., 2008. The probiotic Escherichia coli strain Nissle 1917 induces gamma delta T cell apoptosis via caspase- and FasL-dependent pathways 20, 829–840. https://doi.org/10.1093/intimm/dxn041

Hafez, M., Hayes, K., Goldrick, M., Grencis, R.K., Roberts, I.S., 2010. The K5 capsule of Escherichia coli strain nissle 1917 is important in stimulating expression of toll-like receptor 5, CD14, MyD88, and TRIF together with the induction of interleukin-8 expression via the mitogen-activated protein kinase pathway in epithelial cells. Infect. Immun. 78, 2153–2162. https://doi.org/10.1128/IAI.01406-09

Harden, N., Hau Wang, S.J., Krieger, C., 2016. Making the connection - Shared molecular machinery and evolutionary links underlie the formation and plasticity of occluding junctions and synapses. J. Cell Sci. 129, 3067–3076. https://doi.org/10.1242/jcs.186627

Heine, H., Lien, E., 2003. Toll-like receptors and their function in innate and adaptive immunity. Int. Arch. Allergy Immunol. https://doi.org/10.1159/000069517

Hoffmann, J.A., Kafatos, F.C., Janeway, C.A., Ezekowitz, R.A.B., 1999. Phylogenetic perspectives in innate immunity. Science (80). https://doi.org/10.1126/science.284.5418.1313

Imperial, I.C.V.J., Ibana, J.A., 2016. Addressing the Antibiotic Resistance Problem with Probiotics: Reducing the Risk of Its Double-Edged Sword Effect. Front. Microbiol. 07, 1983. https://doi.org/10.3389/fmicb.2016.01983

Islam, S.U., 2016. Clinical Uses of Probiotics. Medicine (Baltimore). 95, e2658. https://doi.org/10.1097/MD.0000000000002658

Jerome, J.P., Bell, J.A., Plovanich-Jones, A.E., Barrick, J.E., Brown, C.T., Mansfield, L.S., 2011. Standing genetic variation in contingency loci drives the rapid adaptation of Campylobacter jejuni to a novel host. PLoS One 6. https://doi.org/10.1371/journal.pone.0016399

Juge, N., 2012. Microbial adhesins to gastrointestinal mucus. Trends Microbiol. https://doi.org/10.1016/j.tim.2011.10.001

Khan, I., Prakash, A., Agashe, D., 2017. Experimental evolution of insect immune memory versus pathogen resistance. Proc. R. Soc. B Biol. Sci. 284, 20171583. https://doi.org/10.1098/rspb.2017.1583

Kleta, S., Nordhoff, M., Tedin, K., Wieler, L.H., Kolenda, R., Oswald, S., Oelschlaeger, T.A., Bleiß, W., Schierack, P., 2014. Role of F1C Fimbriae, Flagella, and secreted bacterial components in the inhibitory effect of probiotic Escherichia coli Nissle 1917 on atypical enteropathogenic E. coli infection. Infect. Immun. 82, 1801–1812. https://doi.org/10.1128/IAI.01431-13

Kurtz, C., Denney, W.S., Blankstein, L., Guilmain, S.E., Machinani, S., Kotula, J., Saha, S., Miller, P., Brennan, A.M., 2018. Translational Development of Microbiome-Based Therapeutics: Kinetics of E. coli Nissle and Engineered Strains in Humans and Nonhuman Primates. Clin. Transl. Sci. 11, 200–207. https://doi.org/10.1111/cts.12528

Landini, P., Zehnder, A.J.B., 2002. The global regulatory hns gene negatively affects adhesion to solid surfaces by anaerobically grown Escherichia coli by modulating expression of flagellar genes and lipopolysaccharide production. J. Bacteriol. 184, 1522–1529. https://doi.org/10.1128/JB.184.6.1522-1529.2002

Lee, H., Popodi, E., Tang, H., Foster, P.L., 2012. Rate and molecular spectrum of spontaneous mutations in the bacterium Escherichia coli as determined by whole-genome sequencing. Proc. Natl. Acad. Sci. U. S. A. 109, E2774–E2783. https://doi.org/10.1073/pnas.1210309109

Li, C., Louise, C.J., Shi, W., Adler, J., 1993. Adverse conditions which cause lack of flagella in Escherichia coli. J. Bacteriol. 175, 2229–2235. https://doi.org/10.1128/jb.175.8.2229-2235.1993

Lionakis, M.S., 2011. Drosophila and Galleria insect model hosts: New tools for the study of fungal virulence, pharmacology and immunology. Virulence. https://doi.org/10.4161/viru.2.6.18520

Lorenz, A., Schulze, J., 1996. Establishment of E. coli NISSLE 1917 and its interaction with Candida albicans in gnotobiotic rats. Microecol Ther 24, 45–51.

Mandel, L., Trebichavsky, I., Splichal, I., Schulze, J., 1995. Stimulation of intestinal immune cells by E. coli in gnotobiotic piglets, in: Advances in Experimental Medicine and Biology. Springer, Boston, MA, pp. 463–464. https://doi.org/10.1007/978-1-4615-1941-6_96

Milutinović, B., Stolpe, C., Peuß, R., Armitage, S.A.O., Kurtz, J., 2013. The Red Flour Beetle as a Model for Bacterial Oral Infections. PLoS One. https://doi.org/10.1371/journal.pone.0064638

Misof, B., Liu, S., Meusemann, K., Peters, R.S., Donath, A., Mayer, C., Frandsen, P.B., Ware, J., Flouri, T., Beutel, R.G., Niehuis, O., Petersen, M., Izquierdo-Carrasco, F., Wappler, T., Rust, J., Aberer, A.J., Aspöck, U., et al., 2014. Phylogenomics resolves the timing and pattern of insect evolution. Science (80). 346, 763–767. https://doi.org/10.1126/science.1257570

Moghadam, N.N., Thorshauge, P.M., Kristensen, T.N., de Jonge, N., Bahrndorff, S., Kjeldal, H., Nielsen, J.L., 2018. Strong responses of Drosophila melanogaster microbiota to developmental temperature. Fly (Austin). 12, 1–12. https://doi.org/10.1080/19336934.2017.1394558

Nissle, A., 1919. Weiteres über die Mutaflorbehandlung unter besonderer Berücksichtigung der chronischen Ruhr. Münch. Med. Wschr. 678–681.

O’Toole, P.W., Marchesi, J.R., Hill, C., 2017. Next-generation probiotics: The spectrum from probiotics to live biotherapeutics. Nat. Microbiol. https://doi.org/10.1038/nmicrobiol.2017.57

Perini, H.F., Moralez, A.T.P., Almeida, R.S.C., Panagio, L.A., Junior, A.O.G., Barcellos, F.G., Furlaneto-Maia, L., Furlaneto, M.C., 2019. Phenotypic switching in Candida tropicalis alters host-pathogen interactions in a Galleria mellonella infection model. Sci. Rep. 9. https://doi.org/10.1038/s41598-019-49080-6

Pharma-Zentrale GmbH, 2020. Mutaflor® [WWW Document]. URL https://www.mutaflor.com/mutaflor-clinically-proven-efficacy/introduction-and-overview.html

Reister, M., Hoffmeier, K., Krezdorn, N., Rotter, B., Liang, C., Rund, S., Dandekar, T., Sonnenborn, U., Oelschlaeger, T.A., 2014. Complete genome sequence of the gram-negative probiotic escherichia coli strain Nissle 1917. J. Biotechnol. 187, 106–107. https://doi.org/10.1016/j.jbiotec.2014.07.442

Roth, O., Sadd, B.M., Schmid-Hempel, P., Kurtz, J., 2009. Strain-specific priming of resistance in the red flour beetle, *Tribolium castaneum*. Proceedings. Biol. Sci. 276, 145–51. https://doi.org/10.1098/rspb.2008.1157

Roudsari, M.R., Karimi, R., Sohrabvandi, S., Mortazavian, A.M., 2015. Health Effects of Probiotics on the Skin. Crit. Rev. Food Sci. Nutr. 55, 1219–1240. https://doi.org/10.1080/10408398.2012.680078

Sabharwal, H., Cichon, C., Ölschläger, T.A., Sonnenborn, U., Alexander Schmidt, M., 2016. Interleukin-8, CXCL1, and MicroRNA miR-146a responses to probiotic Escherichia coli Nissle 1917 and enteropathogenic E. coli in human intestinal epithelial T84 and monocytic THP-1 cells after apical or basolateral infection. Infect. Immun. 84, 2482–2492. https://doi.org/10.1128/IAI.00402-16

Santacroce, L., Charitos, I.A., Bottalico, L., 2019. A successful history: probiotics and their potential as antimicrobials. Expert Rev. Anti. Infect. Ther. https://doi.org/10.1080/14787210.2019.1645597

Sassone-Corsi, M., Nuccio, S.P., Liu, H., Hernandez, D., Vu, C.T., Takahashi, A.A., Edwards, R.A., Raffatellu, M., 2016. Microcins mediate competition among Enterobacteriaceae in the inflamed gut. Nature 540, 280–283. https://doi.org/10.1038/nature20557

Schulze, J., Lorenz, A., Mandel, L., 1992. Colonisation of Escherichia coli in different gnotobiotic animal models. Microb. Ecol Heal. Dis 5, 4–5.

Scully, L.R., Bidochka, M.J., 2006. Developing insect models for the study of current and emerging human pathogens. FEMS Microbiol. Lett. 263, 1–9. https://doi.org/10.1111/j.1574-6968.2006.00388.x

Sokoloff, A., 1977. The biology of Tribolium with special emphasis on genetic aspects. Volume 3.

Somerville, G.A., Beres, S.B., Fitzgerald, J.R., DeLeo, F.R., Cole, R.L., Hoff, J.S., Musser, J.M., 2002. In vitro serial passage of Staphylococcus aureus: Changes in physiology, virulence factor production, and agr nucleotide sequence. J. Bacteriol. 184, 1430–1437. https://doi.org/10.1128/JB.184.5.1430-1437.2002

Soutourina, O.A., Bertin, P.N., 2003. Regulation cascade of flagellar expression in Gram-negative bacteria. FEMS Microbiol. Rev. https://doi.org/10.1016/S0168-6445(03)00064-0

Soutourina, O.A., Krin, E., Laurent-Winter, C., Hommais, F., Danchin, A., Bertin, P.N., 2002. Regulation of bacterial motility in response to low pH in Escherichia coli: The role of H-NS protein. Microbiology 148, 1543–1551. https://doi.org/10.1099/00221287-148-5-1543

Souza, É.L., Elian, S.D., Paula, L.M., Garcia, C.C., Vieira, A.T., Teixeira, M.M., Arantes, R.M., Nicoli, J.R., Martins, F.S., 2016. Escherichia coli strain Nissle 1917 ameliorates experimental colitis by modulating intestinal permeability, the inflammatory response and clinical signs in a faecal transplantation model. J. Med. Microbiol. 65, 201–210. https://doi.org/10.1099/jmm.0.000222

Sturm, A., Rilling, K., Baumgart, D.C., Gargas, K., Abou-Ghazalé, T., Raupach, B., Eckert, J., Schumann, R.R., Enders, C., Sonnenborn, U., Wiedenmann, B., Dignass, A.U., 2005. Escherichia coli Nissle 1917 distinctively modulates T-cell cycling and expansion via Toll-like receptor 2 signaling. Infect. Immun. 73, 1452–1465. https://doi.org/10.1128/IAI.73.3.1452-1465.2005

Suzuki, Y., Truman, J.W., Riddiford, L.M., 2008. The role of broad in the development of Tribolium castaneum: Implications for the evolution of the holometabolous insect pupa. Development 135, 569–577. https://doi.org/10.1242/dev.015263

Tate, A.T., Andolfatto, P., Demuth, J.P., Graham, A.L., 2017. The within-host dynamics of infection in trans-generationally primed flour beetles. Mol. Ecol. 26, 3794–3807. https://doi.org/10.1111/mec.14088

Thomas, C.M., Versalovic, J., 2010. Probiotics-host communication. Gut Microbes 1, 148–163. https://doi.org/10.4161/gmic.1.3.11712

Tomoyasu, Y., Denell, R.E., 2004. Larval RNAi in Tribolium (Coleoptera) for analyzing adult development. Dev. Genes Evol. 214, 575–578. https://doi.org/10.1007/s00427-004-0434-0

Troge, A., Scheppach, W., Schroeder, B.O., Rund, S.A., Heuner, K., Wehkamp, J., Stange, E.F., Oelschlaeger, T.A., 2012. More than a marine propeller - the flagellum of the probiotic Escherichia coli strain Nissle 1917 is the major adhesin mediating binding to human mucus. Int. J. Med. Microbiol. 302, 304–314. https://doi.org/10.1016/j.ijmm.2012.09.004

Ukena, S.N., Singh, A., Dringenberg, U., Engelhardt, R., Seidler, U., Hansen, W., Bleich, A., Bruder, D., Franzke, A., Rogler, G., Suerbaum, S., Buer, J., Gunzer, F., Westendorf, A.M., 2007. Probiotic Escherichia coli Nissle 1917 inhibits leaky gut by enhancing mucosal integrity. PLoS One 2. https://doi.org/10.1371/journal.pone.0001308

Vassiliadis, G., Destoumieux-Garzón, D., Lombard, C., Rebuffat, S., Peduzzi, J., 2010. Isolation and characterization of two members of the siderophore-microcin family, microcins M and H47. Antimicrob. Agents Chemother. 54, 288–297. https://doi.org/10.1128/AAC.00744-09

Wehkamp, J., Harder, J., Wehkamp, K., Wehkamp-Von Meissner, B., Schlee, M., Enders, C., Sonnenborn, U., Nuding, S., Bengmark, S., Fellermann, K., Schröder, J.M., Stange, E.F., 2004. NF-κB- and AP-1-mediated induction of human beta defensin-2 in intestinal epithelial cells by Escherichia coli Nissle 1917: A novel effect of a probiotic bacterium. Infect. Immun. 72, 5750–5758. https://doi.org/10.1128/IAI.72.10.5750-5758.2004

Weiss, G., 2013. Intestinal irony: How probiotic bacteria outcompete bad bugs. Cell Host Microbe. https://doi.org/10.1016/j.chom.2013.07.003

Wielgoss, S., Schneider, D., Barrick, J.E., Tenaillon, O., Cruveiller, S., Chane-Woon-Ming, B., Médigue, C., Lenski, R.E., 2011. Mutation rate inferred from synonymous substitutions in a long-term evolution experiment with escherichia coli. G3 Genes, Genomes, Genet. 1, 183–186. https://doi.org/10.1534/g3.111.000406

Willott, E., Balda, M.S., Fanning, A.S., Jameson, B., Van Itallie, C., Anderson, J.M., 1993. The tight junction protein ZO-1 is homologous to the Drosophila discs-large tumor suppressor protein of septate junctions. Proc. Natl. Acad. Sci. U. S. A. 90, 7834–7838. https://doi.org/10.1073/pnas.90.16.7834

Wittchen, E.S., Haskins, J., Stevenson, B.R., 1999. Protein interactions at the tight junction. Actin has multiple binding partners, and ZO-1 forms independent complexes with ZO-2 and ZO-3. J. Biol. Chem. 274, 35179–35185. https://doi.org/10.1074/jbc.274.49.35179

Yokoi, K., Koyama, H., Minakuchi, C., Tanaka, T., Miura, K., 2012. Antimicrobial peptide gene induction, involvement of Toll and IMD pathways and defense against bacteria in the red flour beetle, Tribolium castaneum. Results Immunol. 2, 72–82. https://doi.org/10.1016/j.rinim.2012.03.002

Zogaj, X., Nimtz, M., Rohde, M., Bokranz, W., Romling, U., 2001. The multicellular morphotypes of Salmonella typhimurium and Escherichia coli produce cellulose as the second component of the extracellular matrix. Mol. Microbiol. 39, 1452–1463. https://doi.org/10.1046/j.1365-2958.2001.02337.x

Zou, Z., Evans, J.D., Lu, Z., Zhao, P., Williams, M., Sumathipala, N., Hetru, C., Hultmark, D., Jiang, H., 2007. Comparative genomic analysis of the Tribolium immune system. Genome Biol. 8, R177. https://doi.org/10.1186/gb-2007-8-8-r177

Zyrek, A.A., Cichon, C., Helms, S., Enders, C., Sonnenborn, U., Schmidt, M.A., 2007. Molecular mechanisms underlying the probiotic effects of Escherichia coli Nissle 1917 involve ZO-2 and PKCζ redistribution resulting in tight junction and epithelial barrier repair. Cell. Microbiol. 9, 804–816. https://doi.org/10.1111/j.1462-5822.2006.00836.x

