## Supplemental information for "Serial passage of the human probiotic *E. coli* Nissle 1917 in an insect host leads to changed bacterial phenotypes"

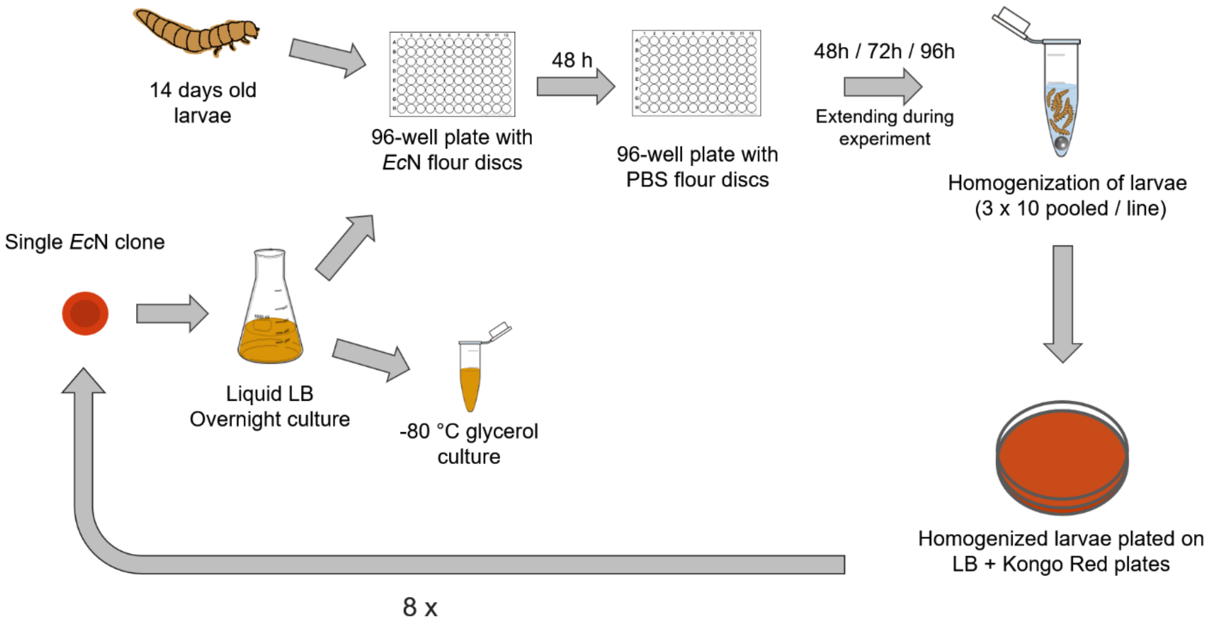


Figure S1: Serial passage workflow: 48 larvae were used per replicate line per passage.

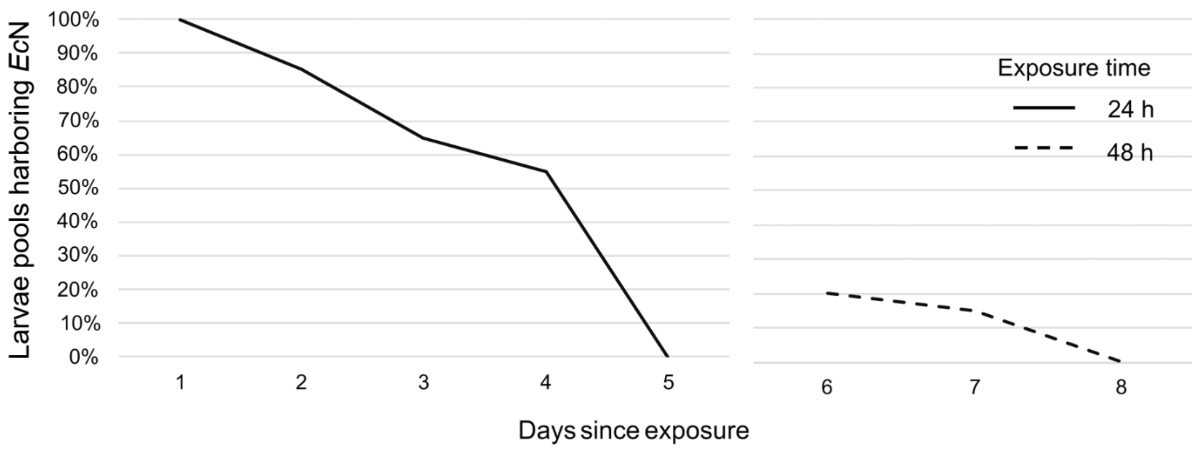


Figure S2: Persistence of *Ec*N in *T. castaneum* larvae. The proportion of larvae harboring *Ec*N after 24 h and 48 h of oral exposure. Per day 20 pools of 10 larvae were tested for bacterial abundance (n = 160).

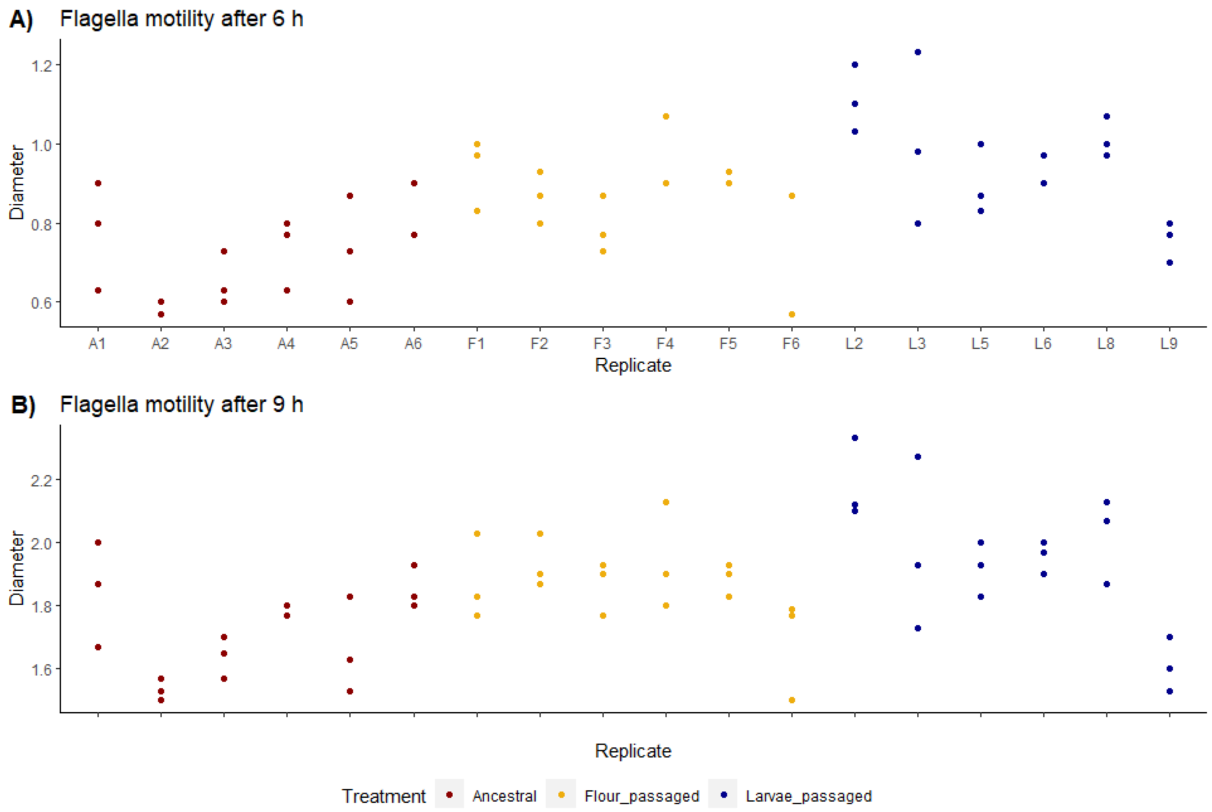


Figure S3: Differences in swarming ability between the individual replicates of the evolved lines and the ancestral strain. Triplicates of six replicates per treatment were analyzed. The radius of the swarmed areas on 0.3 % agar plates was measured after 6 h (A) and 9 h (3B) post inoculation.

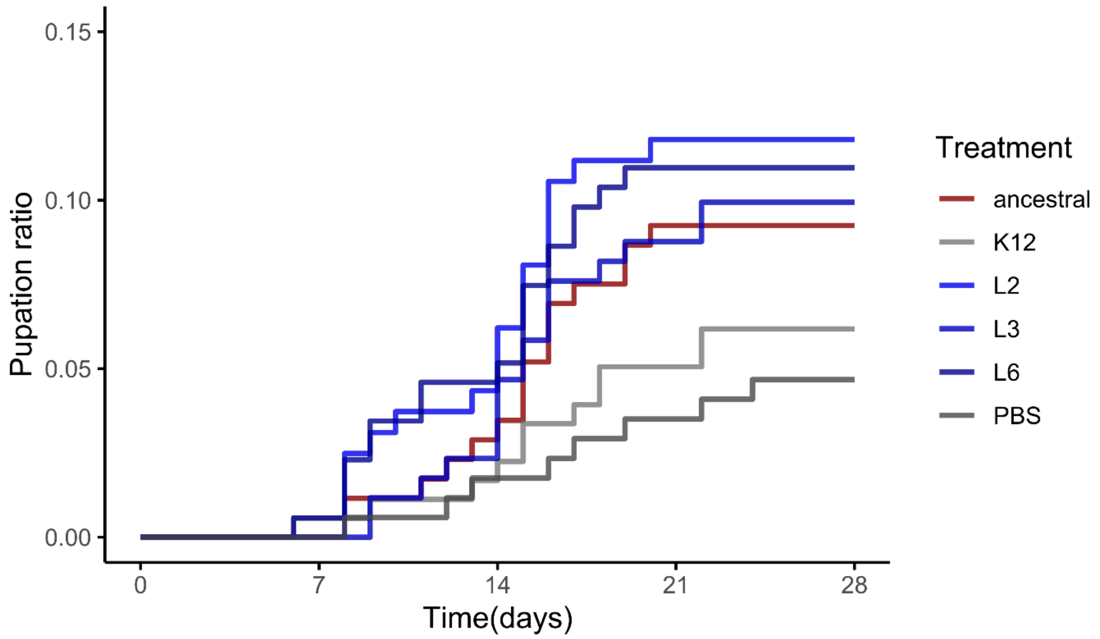


Figure S4: *T. castaneum* pupation upon *E. coli* treatment. The proportion of pupated beetle larvae (14 days old) exposed to *E. coli*-containing flour diet for 72 h (5.3 x 1010 cells / g flour), before transfer to PBS. Three passaged *Ec*N strains (L2, L3, L6), as well as the ancestral strain, were used for pretreatment. PBS served as a negative control for pretreatment and treatment. The larvae were individualized in 96-well plates (n = 1028).

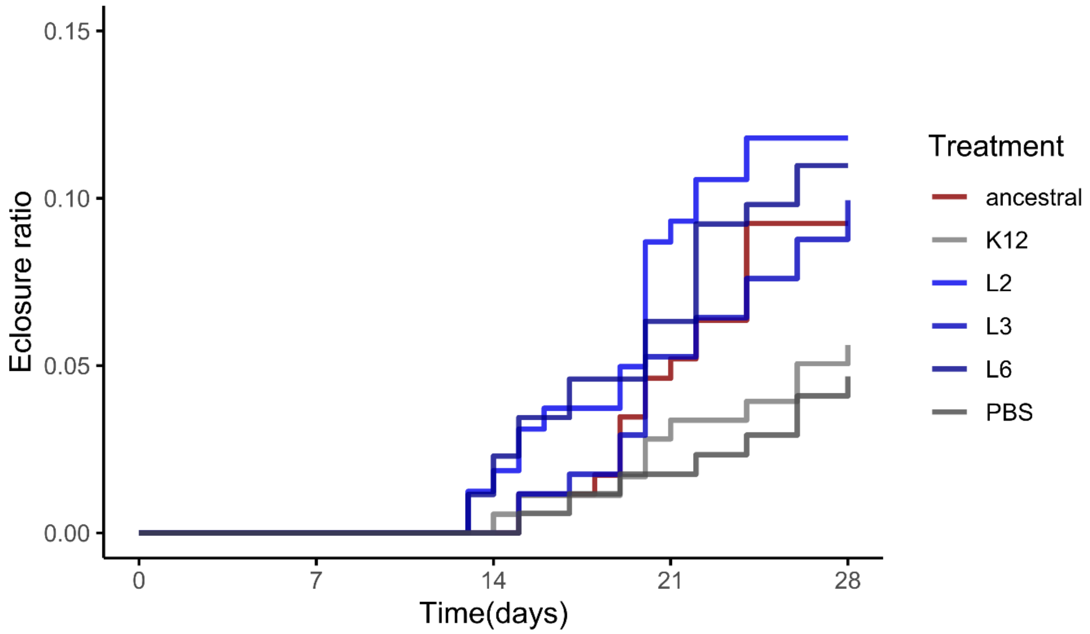


Figure S5: *T. castaneum* eclosure upon *E. coli* treatment. The proportion of beetle larvae (14 days old) exposed to *E. coli*-containing flour diet for 72 h (5.3 x 1010 cells / g flour), before transfer to PBS. Three passaged *Ec*N strains (L2, L3, L6), as well as the ancestral strain, were used for pretreatment. PBS served as a negative control for pretreatment and treatment. The larvae were individualized in 96-well plates (n = 1028).

Transparent methods

### 1 Model organisms

#### *1.1 Tribolium castaneum*

All experiments in this work involving red flour beetles were conducted with the *Tribolium castaneum* strain Cro1, which was collected from a granary in Croatia in Summer 2010 (Milutinović *et al.*, 2013). The beetles were kept on heat sterilized (75 °C; overnight) all-purpose wheat flour (Type 550) with 5 wt.% yeast at 30 °C and 70 % air humidity. The incubators were set up for a 12h/12h light-dark cycle. Synchronization of age was achieved by transferring ~ 1-month-old adult beetles to a cultivation box containing fresh flour with yeast. After 24 h of oviposition, the adults were removed by sieving the flour with a metal sieve and a mesh size of 720 µm (Retsch).

#### *1.2 Escherichia coli* Nissle 1917

The *Escherichia coli* Nissle 1917 (*Ec*N) strain used for the serial passage experiment was obtained from Ardeypharm GmbH (Herdecke) as the probiotic product Mutaflor® and kept in the laboratory of U. Dobrindt (Institute of Hygiene, University of Münster). For long term storage, glycerol was added to an LB (lysogeny broth) overnight culture (25% final glycerol concentration). It was stored in 500 µL microtubes as 200 µL aliquots of 100 µL at - 80 °C. Single colonies for lab use of *Ec*N were generated by streaking out the -80 °C glycerol culture on LB agar plates and incubating at 30 °C overnight. Adding a 40 µg/mL Congo Red and 20 µg/mL Coomassie Brilliant Blue the LB agar medium produces red agar plates that are used for the characterization of *Ec*N colonies. *Ec*N forms characteristic red dry and rough colonies on this agar, by incorporating the dyes in the colonies' biofilm. This mechanism is used to selectively identify *Ec*N colonies.

#### 1.3 *Bacillus thuringiensis* bv. *tenebrionis*

The *Bacillus thuringiensis* bv. *tenebrionis* (*Btt*, Krieg *et al.*, 2009) strain used in this work was acquired from the *Bacillus* Genetic Stock Center of the Ohio State University (USA) and was stored in 500 µL microtubes as 100 µL aliquots with 25 vol.% glycerol. Single *Btt* colonies were generated by streaking out the -80 °C glycerol culture on LB agar plates and incubating at 30 °C overnight.

### 2 Serial passage

#### 2.1 Experimental design

Eight oral uptake cycles were performed by introducing *Ec*N to the environment of the larval midgut using oral exposure.

The experiment consisted of 10 replicate lines passaged through larvae and 6 replicate lines passaged through flour. It was conducted in two equally sized blocks. All lines were initially derived from one ancestral *Ec*N strain (Figure S1)

2.2 Liquid cultures

Initially, frozen ancestral *Ec*N stock cultures and later *Ec*N cells extracted from larvae were plated on LB agar plates dyed with Congo Red and Coomassie Blue and incubated overnight at 30 °C. The following day, 50 mL of liquid LB were inoculated with a single colony and incubated at 30 °C and 180 rpm overnight. After centrifugation and washing the cultures with PBS at 4000 g at room temperature (RT) for 10 minutes, the cell concentrations were standardized to 8 x $10^9$ cells/mL using an approximation based on the optical density (turbidity) of the sample, which was measured at a wavelength of 600 nm ($OD_{600}$). The $OD_{600}$-based calculations were calibrated using an *Ec*N-specific standard curve (produced previously, data not shown), which was verified by streaking out several dilutions of measured samples and counting colony-forming units (CFUs). Aliquots of cell suspensions with adjusted concentrations were stored as frozen glycerol cultures for each line and passage as backups in case a passage needed to be restarted. For this, 100 µL of cell suspension were added to 100 µL of 50 % glycerol in 500 µL microtubes and stored in a -80 °C freezer.

*2.3 Ec*N uptake

*Ec*N bacteria were fed individually to 14-day-old larvae according to an oral exposure protocol modified from Milutinović *et al.*, 2013. The standardized overnight liquid bacterial cell (2.2.) solutions of the ancestral and the 10 passaged lines were therefore mixed with flour and pipetted to half of a 96-well plate per line with a volume of 20 µL / well. With a concentration of 5.3 x $10^{10}$ cells / g flour, each of the resulting 'flour discs' contained about 1.6 x $10^8$ cells in total. The discs were left to dry overnight at 30°C. Per replicate line, 48 14 day-old larvae were exposed to the bacteria upon transferring larvae into these 96-well plates with the flour discs containing EcN . After 24h of continuous exposure to *Ec*N at 30 °C and 70 % humidity, individual larvae were transferred to 96-well plates containing flour discs without *Ec*N. We thereby avoided direct transfer of EcN and extracted only bacteria that had persisted in the host, as described in the following chapter.

2.4 Re-isolation of *Ec*N

The bacteria were extracted at extending time points to maintain a selection pressure. In passages, 1 to 4 *Ec*N was extracted after 48 h being on the discs without *Ec*N, in passages 5 to 7 after 72 h, and in passage 8 after 96 h.

Before the extraction of bacteria, all larvae were cleaned and surface sterilized with three steps. First by dipping into sterile water for 10 seconds for washing residual flour, followed by dipping for 10 seconds into 70% ethanol, and dipping into sterile water for 10 seconds to remove residual ethanol. Three pools of 10 surface-sterilized larvae per line were transferred to 500 µL microtubes containing 200 µL PBS and a sterile metal bead. The larvae were homogenized using a Mixer Mill MM301 (Retsch) for 2 minutes at 28 Hz. The homogenates were then plated out on LB agar plates dyed with Congo Red and Coomassie Blue to allow the identification of red *Ec*N colonies, which incorporate the dyes in their biofilm. After 24 h of incubation at 30 °C, one colony per line was used to inoculate a new 50 mL overnight culture starting the next passage.

2.5 Flour disc passage

To test for potential adaptation of *Ec*N to the flour environment, 6 flour-passaged lines were treated similarly as the larvae-passaged lines, but passaged only through flour discs (at 30 °C and 70 % humidity) that did not contain any larvae. In parallel to the extraction of EcN from larvae in the larvae-passaged lines, the flour discs with the flour-passaged lines were resuspended in 200 µL of PBS and streaked out on LB agar plates dyed with Congo Red and Coomassie Blue, using an inoculation loop. A single colony per control line was used to inoculate the liquid culture for the next passage.

### 3 Phenotypic analyses

3.1 Persistence analysis

To analyze the persistence of the serially passaged *Ec*N in the *T. castaneum* host, an infection experiment was performed that included all larvae-passaged *Ec*N lines after 8 passages, as well as the ancestral strain. For this, per *Ec*N line, 144 larvae (i.e. 1584 larvae in total) were orally exposed to vegetative bacterial cells for 48 h. In contrast to the preceding passages, in which exposed larvae were sampled in pools of 10 larvae only once per passage cycle (see 2.4), we now monitored persistence in detail, i.e. in daily intervals by checking the presence of *Ec*N in 20 individual larvae per day and line. Droplets of 10 µL of 10 individual homogenates were applied per LB + Congo Red / Coomassie Blue plate. After overnight incubation at 30 °C, the plates were screened for the presence or absence of *Ec*N colonies.

3.2 Growth dynamics

Possible differences in the growth dynamics of the passaged replicates were assessed by monitoring the turbidity of liquid cultures over time. Liquid LB medium cultures with a volume of 5 mL were inoculated with 6 larvae passaged replicates (L2, L3, L5, L6, L8, L9), 6 flour passaged replicates (F1-6), and 6 pseudoreplicates of the ancestral strain. Pseudoreplicates of ancestral strain were produced by inculating 6 clones in 6 different cultures. The samples were incubated overnight at 30 °C and 180 rpm. All cultures were diluted to a relative absorbance of 0.5 at a wavelength of 600 nm ($OD_{600}$). Of the standardized cell solutions, 10 µL were used to inoculate 200 µL of liquid LB medium (diluted 1:10 with PBS) in a 96-well plate. For each bacterial line, 4 technical replicates were measured every 15 min for 24 h at 30 °C using the Infinite® 200 PRO plate reader (Tecan).

3.3 Motility

The swimming motility of 3 technical replicates of 6 biological replicates per passage treatment (L2, L3, L5, L6, L8, L9, F1-6) and 6 pseudo-repicates of the ancestral strain was tested. LB agar plates with 0.3 % agar can be used to evaluate the motility of bacterial strains (Arora et al., 1998). Bacterial cells were transferred from single colonies to the center of an LB 0.3 % agar plate by slightly touching the agar surface, using the tip of a sterile toothpick. The plates were incubated at 30 °C and the diameter of the swimming area was measured after 6 h and 9 h.

3.4. Colony morphology

The synthesis of cellulose and curli fimbriae, components of the extracellular matrix, can be visualized using the dyes Calcofluor white (under UV) and Congo Red, respectively (Cimdins and Simm, 2017; Zogaj et al., 2001). To test the colony morphology and the formation of an extracellular matrix, bacterial strains were streaked out on agar plates with added Congo Red or Calcoflour white and incubated at 30 °C and 37 °C for 96 h.

### 4. Whole-genome sequencing analysis

#### 4.1. DNA extraction and whole-genome sequencing

The genomes of the three passaged lines L2, F4 and L9 were sequenced using next-generation sequencing. While L2 and F4 were chosen based on a trend for elevated persistence, L9 was put forward for sequencing because of its peculiar colony morphology.The total genomic DNA was isolated using the MagAttract® HMW DNA kit (Qiagen, Hilden, Germany). To prepare 500 bp paired-end libraries of all isolates we used the Nextera XT DNA Library Preparation kit (Illumina, San Diego, CA, USA). Libraries were sequenced on the Illumina MiSeq sequencing platform using v2 sequencing chemistry.

#### 4.2. Quality control and variant calling

The quality of the raw sequencing data was analyzed using FastQC v0.11.5 (Andrews, 2010). Raw reads were trimmed using Sickle v1.33 (https://github.com/najoshi/sickle). Quality trimmed reads were then aligned against the in-house reference *Ec*N genome (CP058217) using Burrows-Wheeler Aligner (BWA) v0.7.17 (Li and Durbin, 2010). The produced alignment was sorted, and duplicates were marked using Picard v2.17.3 (https://github.com/broadinstitute/picard). Variant calling and filtering were performed using the Genome Analysis Toolkit (GATK) v3.8.0 (McKenna et al., 2010). The effects of the resulting variants were then annotated using SnpEff v4.3 (Cingolani et al., 2012).

#### 4.3. *De novo*-assembly and pan-genome analysis

Genome assembly of the processed reads was carried out with SPAdes v3.13.1 (Bankevich et al., 2012). The assembled genomes were then annotated using Prokka v1.12 (Seemann, 2014) and used for a pan-genome analysis, which was performed using Roary v3.12.0 (Page et al., 2015). The resulting presence-absence matrix of orthologous genes was visualized using FriPan (http://drpowell.github.io/FriPan/).

### 5. Effects of *Ec*N on host mortality and gene expression

#### 5.1 Oral exposure with *Ec*N and *Btt*

To investigate the putative protective effect of *Ec*N, larvae were pre-treated with the passaged *Ec*N strains L2, L3 and L6 as well as with the ancestral strain (5.3 x $10^{10}$ cells / g flour) and subsequently exposed to *Btt* spores ( 3.3 x $10^{10}$ spores / g flour).

Mortality was screened for 14 days and pupation/eclosure for 28 days. Dead larvae were identified by immobility, the characteristic body shape, and a darkened color.

*Ec*N cells were fed to 14 day-old larvae according to the modified oral exposure protocol described in Milutinović et al., 2013. (see chapter 2.2. and 2.3.). After 24 h on the *Ec*N diet, the larvae were transferred to the *Btt* diet.

The diet for oral exposure with *Btt* was made as described in Milutinović et al., 2013. In short, *Btt* from the frozen stock was plated on an LB agar plate and incubated overnight. The following day, 5 mL of BT medium [w/V–0.75% Bacto Peptone (Sigma), 0.1% glucose, 0.34% KH2PO4, 0.435% K2HPO4] was supplemented with 25 µL of sterile salt solution (0.2 M MgSO4, 2 mM MnSO4, 17 mM ZnSO4, 26 mM FeSO4) and 6.25 µL of sterile 1 M CaCl2 × 2H2O solution, inoculated with 5 single *Btt* colonies from the LB agar plate and incubated overnight in culture tubes (Simport) at 200 rpm. The following morning, 300 mL of BT medium were supplemented with 1.5 mL of salt solution, 375 µL 1M CaCl2 × 2H2O, and inoculated with 5 mL of the overnight culture. The culture was incubated in a 2 L Erlenmeyer flask for 7 days at 180 rpm. After 7 days of sporulation, the spores were centrifuged at 4500 rpm for 15 min at room temperature (RT), followed by washing with phosphate-buffered saline (PBS) and centrifuged again. *Btt* spores were counted using a Thoma counting chamber. The adjusted concentration of liquid bacterial cultures was mixed with flour and pipetted into 96-well plates. After drying the plates, 14 day-old larvae were individually transferred to each well and monitored for survival and development.

#### 5.2. Gene expression upon oral uptake of EcN

To characterize a possible differential immune reaction of *T. castaneum* to the exposure to *Ec*N, the expression levels of several defense genes were assessed. These genes are either involved in the regulation of antimicrobial peptide (AMP) expression or code for AMPs themselves. After oral exposure to *Ec*N, *E. coli* K-12 MG1655 (K12), or a PBS negative control for 3 days, 19 larvae per treatment were snap-frozen in individual microtubes using liquid nitrogen. For the RT-qPCR we used 7 genes, including two housekeeping genes Rp49 and Rpl13a as in Eggert et al., 2014. (Table 1). We used 6 replicates per bacterial treatment.

#### 5.3. Quantitative reverse transcription PCR

Total RNA from 6 samples of 5 pooled frozen larvae for each treatment (*Ec*N, K-12, PBS) was extracted using a combined protocol (Eggert et al., 2014) of TriFast™ (VWR) and spin columns of the SV Total RNA Isolation System (Promega). The quantity and quality of extracted RNA were evaluated using a NanoPhotometer® P-300 (Implen). Per sample 300 ng of RNA were used for cDNA synthesis with the RevertAid First Strand cDNA Synthesis Kit (Thermo Scientific). The reverse transcription was performed according to the manual of the kit. The quantitative reverse transcription PCR (RT-qPCR) was run on a LightCycler® 480 (Roche®) with SYBR® Green fluorescent dye (Thermo Fischer Scientific). The relative expression of the AMPs Attacin2 (Att2), Cecropin2 (Cec2), Defensin2 (Def2), Defensin3 (Def3), and an Osiris16-like protein was assessed using the housekeeping genes of the ribosomal protein L13a (Rpl13a) and ribosomal protein 49 (Rp49). Technical duplicates were set up for each sample.

Table 1. Sequences of primers used for RT- qPCR

| Protein name | Forward primer sequence (5′–3′) | Reverse primer sequence (5′–3′) | Source |
|---|---|---|---|
| Att2 | CAAACGACCAAAGGGAAACTAAA | TGAACTTGTCCAGTTGCATCGA | Yokoi et al., 2012) |
| Cec2 | GCCGAAGGAGCTGGAAGATTA | TGGTGGTGGAGGTTGTTGGTA | Yokoi et al., 2012 |
| Def2 | CCCTTTTCTGCATCTTCGAAAC | CACATGCGGAATGGTTTAGCT | Yokoi et al., 2012 |
| Def3 | TGCAATCACTGCTTACCCACTT | ACAAGCAGCATGATTCACTTTGA | Yokoi et al., 2012 |
| Rp49 | TTATGGCAAACTCAAACGCAAC | GGTAGCATGTGCTTCGTTTTG | Eggert et al.,2014 |
| Rpl13a | GGCCGCAAGTTCTGTCAC | GGTGAATGGAGCCACTTGTT | Eggert et.al, 2014 |
| Osiris 16 | CGACAAGCCTACTCCC | TGTAGTCGTCCTCCTCGTTC | Lindeza and Barth, 2020 unpublished data. |

6. Statistics

All statistical analyses and figures were performed and produced in R (R Core Team, 2020) using RStudio (RStudio Team, 2020). For producing the plots, the packages ggplot2 (Wickham, 2016) and ggpubr (Alboukadel Kassambara, 2020) were used.

To test for a normal distribution of the analyzed data, a Shapiro–Wilk test was performed (Shapiro and Wilk, 1965). Based on the normality test a parametric or non-parametric test was conducted to assess significance.

Persistence data was analyzed using a binomial generalized linear model (Bates et al., 2015; Venables & Ripley, 2002) using package “lme4” (Bates et al., 2015).

Growth curves were analysed using the package growthcurver (Sprouffske and Wagner, 2016). Furthermore, we analyzed growth rate (r) and carrying capacity (K) with Kruskal- Walllis test (Kruskal and Wallis, 1952) followed by the non-parametric Wilcoxon signed-rank test (Wilcoxon, 1992). corrected for multiple testing using the Benjamini-Hochberg procedure(Benjamini and Hochberg, 1995).

Since the bacterial motility and AMP gene data and datasets were normally distributed, and assumptions were met, one-way ANOVA was performed. The means were compared using Tukey Honest Significant Differences (HSD) (Miller, 1981; Yandell, 1997).

For the analysis of survival, a Cox Proportional Hazards Model was applied with one random effect (Cox, 1972; Ripatti and Palmgren, 2000; Therneau et al., 2003) using coxph function from the “survival” package (Andersen and Gill, 1982; Therneau and Grambsch, 2000). The treatment was defined as the fixed factor, while a putative plate effect was defined as a random factor.The assumption were met and and after fitting the model, the variance between treatments was assessed using a one-way analysis of variance. The means were compared using Tukey's test for post-hoc analysis (Hothorn et al., 2008; Tukey, 1949) with Benjamini-Hochberg adjusted p-values.

Pupation and eclosure data were additionally analyzed using a binomial generalized linear model (Bates et al., 2015; Venables and Ripley, 2002).

### References

Alboukadel Kassambara, 2020. ggpubr: “ggplot2” Based Publication Ready Plots.

Andersen, P.K., Gill, R.D., 1982. Cox’s Regression Model for Counting Processes: A Large Sample Study. Ann. Stat. https://doi.org/10.1214/aos/1176345976

Andrews, S., 2010. FastQC: a quality control tool for high throughput sequence data.

Arora, S.K., Ritchings, B.W., Almira, E.C., Lory, S., Ramphal, R., 1998. The Pseudomonas aeruginosa flagellar cap protein, FliD, is responsible for mucin adhesion. Infect. Immun. 66, 1000–1007. https://doi.org/10.1128/iai.66.3.1000-1007.1998

Bankevich, A., Nurk, S., Antipov, D., Gurevich, A.A., Dvorkin, M., Kulikov, A.S., Lesin, V.M., Nikolenko, S.I., Pham, S., Prjibelski, A.D., Pyshkin, A. V., Sirotkin, A. V., Vyahhi, N., Tesler, G., Alekseyev, M.A., Pevzner, P.A., 2012. SPAdes: A new genome assembly algorithm and its applications to single-cell sequencing. J. Comput. Biol. 19, 455–477. https://doi.org/10.1089/cmb.2012.0021

Bates, D., Mächler, M., Bolker, B.M., Walker, S.C., 2015. Fitting linear mixed-effects models using lme4. J. Stat. Softw. 67, 1–48. https://doi.org/10.18637/jss.v067.i01

Benjamini, Y., Hochberg, Y., 1995. Controlling the False Discovery Rate: A Practical and Powerful Approach to Multiple Testing. J. R. Stat. Soc. Ser. B 57, 289–300. https://doi.org/10.1111/j.2517-6161.1995.tb02031.x

Cimdins, A., Simm, R., 2017. Semiquantitative analysis of the red, dry, and rough colony morphology of salmonella enterica serovar typhimurium and Escherichia coli using congo red, in: Methods in Molecular Biology. Humana Press Inc., pp. 225–241. https://doi.org/10.1007/978-1-4939-7240-1_18

Cingolani, P., Platts, A., Wang, L.L., Coon, M., Nguyen, T., Wang, L., Land, S.J., Lu, X., Ruden, D.M., 2012. A program for annotating and predicting the effects of single nucleotide polymorphisms, SnpEff: SNPs in the genome of Drosophila melanogaster strain w1118; iso-2; iso-3. Fly (Austin). 6, 80–92. https://doi.org/10.4161/fly.19695

Cox, D.R., 1972. Regression Models and Life-Tables. J. R. Stat. Soc. Ser. B 34, 187–220. https://doi.org/10.2307/2985181

Eggert, H., Kurtz, J., Diddens-de Buhr, M.F., 2014. Different effects of paternal trans-generational immune priming on survival and immunity in step and genetic offspring. Proc. R. Soc. B Biol. Sci. 281, 20142089. https://doi.org/10.1098/rspb.2014.2089

Krieg, A., Huger, A.M., Langenbruch, G.A., Schnetter, W., 2009. Bacillus thuringiensis var. tenebrionis: ein neuer, gegenüber Larven von Coleopteren wirksamer Pathotyp. Zeitschrift für Angew. Entomol. 96, 500–508. https://doi.org/10.1111/j.1439-0418.1983.tb03704.x

Kruskal, W.H., Wallis, W.A., 1952. Use of Ranks in One-Criterion Variance Analysis. J. Am. Stat. Assoc. 47, 583–621. https://doi.org/10.1080/01621459.1952.10483441

Li, H., Durbin, R., 2010. Fast and accurate long-read alignment with Burrows-Wheeler transform. Bioinformatics 26, 589–595. https://doi.org/10.1093/bioinformatics/btp698

McKenna, A., Hanna, M., Banks, E., Sivachenko, A., Cibulskis, K., Kernytsky, A., Garimella, K., Altshuler, D., Gabriel, S., Daly, M., DePristo, M.A., 2010. The genome analysis toolkit: A MapReduce framework for analyzing next-generation DNA sequencing data. Genome Res. 20, 1297–1303. https://doi.org/10.1101/gr.107524.110

Miller, R.G., 1981. Simultaneous statistical inference. Springer.

Milutinović, B., Stolpe, C., Peuß, R., Armitage, S.A.O., Kurtz, J., 2013. The Red Flour Beetle as a Model for Bacterial Oral Infections. PLoS One. https://doi.org/10.1371/journal.pone.0064638

Page, A.J., Cummins, C.A., Hunt, M., Wong, V.K., Reuter, S., Holden, M.T.G., Fookes, M., Falush, D., Keane, J.A., Parkhill, J., 2015. Roary: Rapid large-scale prokaryote pan genome analysis. Bioinformatics 31, 3691–3693. https://doi.org/10.1093/bioinformatics/btv421

R Core Team, 2020. R: A language and environment for statistical computing.

Ripatti, S., Palmgren, J., 2000. Estimation of multivariate frailty models using penalized partial likelihood. Biometrics 56, 1016–1022. https://doi.org/10.1111/j.0006-341X.2000.01016.x

RStudio Team, 2020. RStudio: Integrated Development Environment for R.

Seemann, T., 2014. Prokka: Rapid prokaryotic genome annotation. Bioinformatics 30, 2068–2069. https://doi.org/10.1093/bioinformatics/btu153

Shapiro, S.S., Wilk, M.B., 1965. An Analysis of Variance Test for Normality (Complete Samples). Biometrika 52, 591. https://doi.org/10.2307/2333709

Sprouffske, K., Wagner, A., 2016. Growthcurver: An R package for obtaining interpretable metrics from microbial growth curves. BMC Bioinformatics 17, 172. https://doi.org/10.1186/s12859-016-1016-7

Therneau, T.M., Grambsch, P.M., 2000. Modeling Survival Data: Extending the Cox Model. Springer.

Therneau, T.M., Grambsch, P.M., Pankratz, V.S., 2003. Penalized survival models and frailty. J. Comput. Graph. Stat. 12, 156–175. https://doi.org/10.1198/1061860031365

Venables, W.N., Ripley, B.D., 2002. Modern Applied Statistics with S. Springer. https://doi.org/10.1007/978-0-387-21706-2_1

Wickham, H., 2016. ggplot2: Elegant Graphics for Data Analysis. Springer-Verlag New York.

Wilcoxon, F., 1992. Individual Comparisons by Ranking Methods, in: Breakthroughs in Statistics. Springer, New York, NY, pp. 196–202. https://doi.org/10.1007/978-1-4612-4380-9_16

Yandell, B.S., 1997. Practical Data Analysis for Designed Experiments. Chapman & Hall.

Yokoi, K., Koyama, H., Minakuchi, C., Tanaka, T., Miura, K., 2012. Antimicrobial peptide gene induction, involvement of Toll and IMD pathways and defense against bacteria in the red flour beetle, Tribolium castaneum. Results Immunol. 2, 72–82. https://doi.org/10.1016/j.rinim.2012.03.002

Zogaj, X., Nimtz, M., Rohde, M., Bokranz, W., Romling, U., 2001. The multicellular morphotypes of Salmonella typhimurium and Escherichia coli produce cellulose as the second component of the extracellular matrix. Mol. Microbiol. 39, 1452–1463. https://doi.org/10.1046/j.1365-2958.2001.02337.x
